# Farewell to single-well: An automated single-molecule FRET platform for high-content, multiwell plate screening of biomolecular conformations and dynamics

**DOI:** 10.1101/2023.02.28.530427

**Authors:** Andreas Hartmann, Koushik Sreenivasa, Mathias Schenkel, Neharika Chamachi, Philipp Schake, Georg Krainer, Michael Schlierf

**Affiliations:** B CUBE Center for Molecular Bioengineering, TU Dresden, Tatzberg 41, 01307 Dresden, Germany; Centre for Misfolding Diseases, Yusuf Hamied Department of Chemistry, University of Cambridge, Lensfield Road, CB2 1EW Cambridge UK; Physics of Life, DFG Cluster of Excellence, TU Dresden, 01062 Dresden, Germany; Department of Bionanoscience, Delft University of Technology, 2629HZ Delft, Netherlands

## Abstract

Single-molecule FRET (smFRET) has become a widely used tool for probing the structure, dynamics, and functional mechanisms of biomolecular systems, and is extensively used to address questions ranging from biomolecular folding to drug discovery. Investigations by smFRET often require sampling of a large parameter space, for example, by varying one or more constituent molecular components in ten or more steps to reliably extract distances, kinetic rates, and other quantitative parameters. Confocal smFRET measurements, for example, which are amongst the widely used smFRET assays, are typically performed in a single-well format and measurements are conducted in a manual manner, making sampling of many experimental parameters laborious and time consuming. To address this challenge, we extend here the capabilities of confocal smFRET beyond single-well measurements by integrating a multiwell plate functionality into a confocal microscope to allow for continuous and automated smFRET measurements. We show that the multiwell plate assay is on par with conventional single-well smFRET measurements in terms of accuracy and precision yet enables probing tens to hundreds of conditions in a fully automized manner. We demonstrate the broad applicability of the multiwell plate assay towards DNA hairpin dynamics, protein folding, and competitive and cooperative protein–DNA interactions, revealing new insights that would be hard if not impossible to achieve with conventional single-well format measurements. The higher sampling density afforded by the multiwell plate format increases the accuracy of data analysis by at least 10-fold. We further showcase that the assay provides access to smFRET-based screening of drug–protein interactions. For the adaptation into existing instrumentations, we provide a detailed guide and open-source acquisition and analysis software. Taken together, the automated multiwell plate assay developed here opens up new possibilities to acquire high-content smFRET datasets for in-depth single-molecule analysis of biomolecular conformations, interactions, and dynamics.

## Introduction

Single-molecule Förster resonance energy transfer (smFRET) has become a widely used technique to monitor biomolecular conformations and dynamics on the nanometer scale^1,2^. Advanced data analysis approaches to characterize dynamics from nanoseconds to minutes and hours, together with standardizations across many laboratories and open science initiatives, have pushed smFRET to a routinely accessible technique in biophysical and biochemical research^3–8^. smFRET, due to its versatility and sensitivity, is now extensively used to address a wide range of questions in dynamic structural biology and biophysics, including the functional mechanisms of enzymes and membrane proteins^9–11^, protein–nucleic acid and small-molecule– protein interactions^12,13^, and protein or nucleic-acid folding^3,14,15^, to name but a few^2,5,16^.

Despite its popularity, smFRET studies often require the curation of large datasets with high statistics and the sampling of a wide parameter space, for example, by varying one or more constituent molecular components in ten or more steps. For instance, structure determination of biomolecular conformations using trilateration approaches requires measurement of multiple directions by FRET, which are often probed in combination with a variation of solution conditions^17,18^. To uncover biomolecular interactions and changes in protein and nucleic-acid structures, for example, in functional investigations or folding studies, ligand or co-solute concentrations are typically varied over several orders of magnitude, giving insights into changes in molecular conformations and kinetics as well as information on binding stoichiometries and affinities^19–22^. Such measurements also often demand a high sampling density to alleviate overparameterization effects and to increase fitting accuracy and precision. Moreover, for the determination of experimental variability, a need for technical and biological replicates arises in general, not least to ascertain reproducibility and enhance scientific rigor^4,8,23^. Hence, rapidly more than 50 conditions need to be probed in smFRET experiments.

Unfortunately, curation of large datasets and sampling of multiple conditions is laborious and time consuming with current smFRET modalities. Confocal smFRET measurements, for example, which are amongst the widely used smFRET assays, are typically performed in a single-well format and measurements are normally conducted in a manual manner, in that, an experimenter needs to replenish and equilibrate the sample after each experiment. Such manual experimentation is also susceptible to changes in environmental conditions, like changes in temperature, instrumental stability, or other parameters. Approaches, based on microfluidic mixing^24^ or multi-spot confocal readouts^25^, have been devised to address these issues, for example, by providing the possibility to vary constituent molecular concentrations or reducing the measurement times per sample chamber. However, these approaches rely on extensive customization and custom-made hardware, are often not easily integratable into existing setups, and typically require expert knowledge not necessarily available in standard lab settings. Moreover, automation is not easily achievable with these approaches.

An attractive, alternative way to address the need for generating large datasets and sampling of a large parameter space are multiwell plates. Multiwell plates are ubiquitous tools in all areas of science because they allow collecting data for tens to hundreds of different conditions.^26^ Hence, implementing multiwell readouts in smFRET experiments should lend itself a powerful approach to probe many different conditions in a fully automated manner within a single continuous experiment under controlled conditions. Fluorescence microscopy experiments for large-scale screening, for example, of single-molecule fluorescence in situ hybridization (smFISH), RNA interference or organoids, have been automated already in many applications based on 96-well or larger multiwell plates^27–29^. However, this powerful platform has not yet been transferred to a format suitable for applications in smFRET experiments.

Here, we introduce an automated smFRET platform for high-content, multiwell plate screening of biomolecular conformations and dynamics. We describe the implementation of multiwell plates for fully automated confocal smFRET measurements in a single, continuous experiment. We provide an open-source software suite for data acquisition, processing, analysis, and visualization. To illustrate the broad applicability of the multiwell plate measurement format, we validate the approach using an array of systems with increasing complexity. Using a DNA ruler system, we show that high precision and accuracy between sample wells can be achieved down to Ångström distances. We further evaluate the possibility to extract millisecond transition rates for DNA nanostructures. In protein-unfolding experiments, we determine thermodynamic stability parameters and parameters related to protein dynamics with high accuracy and sampling density and gain new insights into protein folding mechanisms. We then expand the multiwell plate system to study multi-component systems by probing the competition of two proteins for the same DNA substrate. We thereby discover a new simultaneous binding interaction of RecA and SSB to single-stranded DNA, through the fine sampling in our multiwell plate smFRET assay. Finally, we illustrate the capability to use the multiwell plate smFRET format to screen for small-molecule–protein interactions using a misfolding model of the human cystic fibrosis transmembrane conductance regulator (CFTR) and gain quantitative readouts of the drug–protein interactions. Taken together, we anticipate that our approach will transform smFRET measurements by enabling the acquisition of high-content smFRET datasets for single-molecule analysis in dynamic structural biology and biophysics, and beyond.

## Results and Discussion

### Automated multiwell plate smFRET measurements

Typical commercial or custom-built confocal smFRET instruments allow for individual experiments in a single-well chamber mounted onto a stage for positioning and focusing. Such single-well chambers require cleaning, refilling and equilibration for each experiment, making it laborious for the experimenter to conduct measurements, thereby limiting throughput, and affecting potentially stable conditions for a large set of experiments. Here, we extend the capabilities of confocal smFRET beyond single-well measurements by integrating a multiwell plate functionality into a confocal microscope to allow for continuous and automated smFRET measurements (Fig. 1).

**Figure 1.**
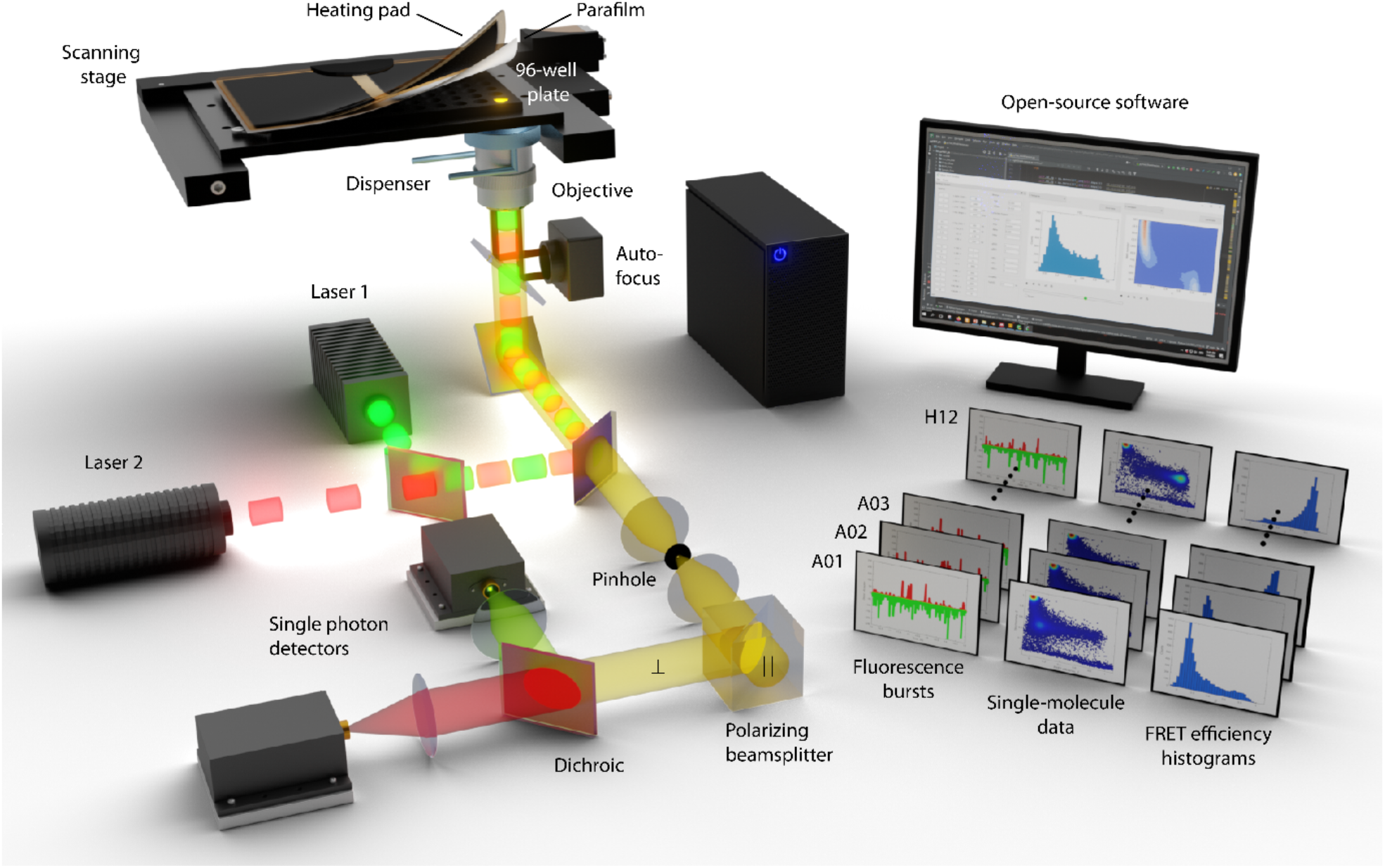
Automated multiwell plate smFRET measurements. Illustration of the automated single-molecule detection setup implementing a multiwell plate functionality for smFRET experiments. A motorized scanning stage holds the multiwell plate, and a script that controls the microscopy stage as well as the data readout proceeds the objective from well to well. During the measurement, a liquid dispenser frequently replaces the immersion medium of the water objective and the autofocus maintains the objective focus at a fixed position in solution. In order to prevent evaporation and condensation, the wells of the plate are sealed, and the top of the well plate is maintained a few degrees Kelvin above the desired temperature by a heating pad. Single-molecule fluorescence recordings are performed using a multi-parameter confocal fluorescence microscope equipped with pulsed lasers, a high-numerical water objective, and single-photon detectors. Photon recordings are done in a spectrally- and polarization-sensitive manner. For simplicity, only the detection pathway of the perpendicular direction (⊥) is shown. A detailed schematic and description of the setup is given in the *SI*. An integrated open-source software suite with graphical user interfaces performs data acquisition, processing, analysis, and visualization. The software is freely available on Github.

To this end, we equipped a multiparameter single-molecule detection microscope with three major additional and commercially available components (Fig. 1, Table S1): (i) a motorized x-y stage, capable of holding a multiwell (*e*.*g*., 96-well) plate with a position accuracy <40 μm; (ii) a heating pad for the multiwell plate to prevent condensation on the well-plate sealing, which is operated a few Kelvin above the desired temperature (*e*.*g*., room temperature); and (iii) a liquid dispenser, which frequently replaces the evaporating immersion medium of the high numerical aperture water objective. During continuous measurements, the confocal detection volume is maintained 30 μm in solution using an autofocus system integrated in the microscope (Table S1).

A custom-written, open-source data acquisition software (Fig. 1), written in Python, synchronizes x-y positioning with data acquisition using predefined libraries of the hardware. Using our software’s graphical user interface (GUI; pyMULTI; available on GitHub), measurement wells and times including descriptive text can be defined. During measurements, the acquired fluorescence data is saved in subfolders for subsequent data analysis. Data analysis is performed by our open-source GUI software (pyBAT and pyVIZ; available on GitHub) and involves fluorescence burst search routines, smFRET analysis, and data visualization, all following state-of-the art analysis protocols^4^. In summary, the new instrument platform with smFRET data acquisition and analysis scripts allows us to perform multiwell plate smFRET experiments in an automated fashion. To test the accuracy and precision of the multiwell plate smFRET platform, we performed 96 independent, but identical smFRET measurements of a mixture of two rigid double-stranded DNA ruler constructs with 9- and 21-base pair (bp) distance between the acceptor and donor fluorophores, respectively (Fig. 2B, Table S4). We loaded the mixture into each well of a 96-well plate, and performed smFRET measurements by taking 20-minutes-long photon recordings of donor and acceptor fluorescence, resulting in a total measurement duration of about 32 h. Subsequently, the data were analyzed using pyBAT and pyVIZ yielding FRET efficiency (*E*) histograms with an average number of ⟨*N*⟩ = 1322 ± 108 bursts per well (Fig. S4). We evaluated the measured *E* values of the two DNA rulers for their accuracy and precision. To this end, we fitted all individual 96 *E*-histograms with two Gaussian distributions to extract the mean FRET efficiencies, the standard deviation, and the ratio of molecules in the 21-bp population (Fig. S4C). Remarkably, we found only minimal deviations between the 96 measurements. The cumulated *E*-histogram of all 96 wells (Fig. 1C, top panel) yielded ⟨*E*_9bp_⟩ = 0.797 and ⟨*E*_21bp_⟩ = 0.146, which agrees very well with the expected FRET efficiencies (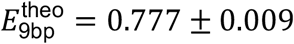 and 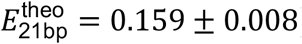) using accessible volume simulations^30^. The mean absolute deviation is < 8.2% (*i*.*e*., 0.8 Å for the 9 bp ruler and 1.1 Å for the 21 bp ruler). We found a very low variation of the measured FRET efficiencies between the individual measurements, as gauged by a standard deviation of *σ*_⟨*E*⟩_ < 0.006. Notably, the minimal loss of molecules throughout the measurement (Fig. S4A) as well as the low variability in the population ratio (Fig. S4D) confirms the measurement stability of the assay. We conclude that our multiwell plate smFRET platform is stable, and we can acquire smFRET data with high accuracy and precision in accordance with conventional single-well sample chamber measurements^4^.

**Figure 2.**
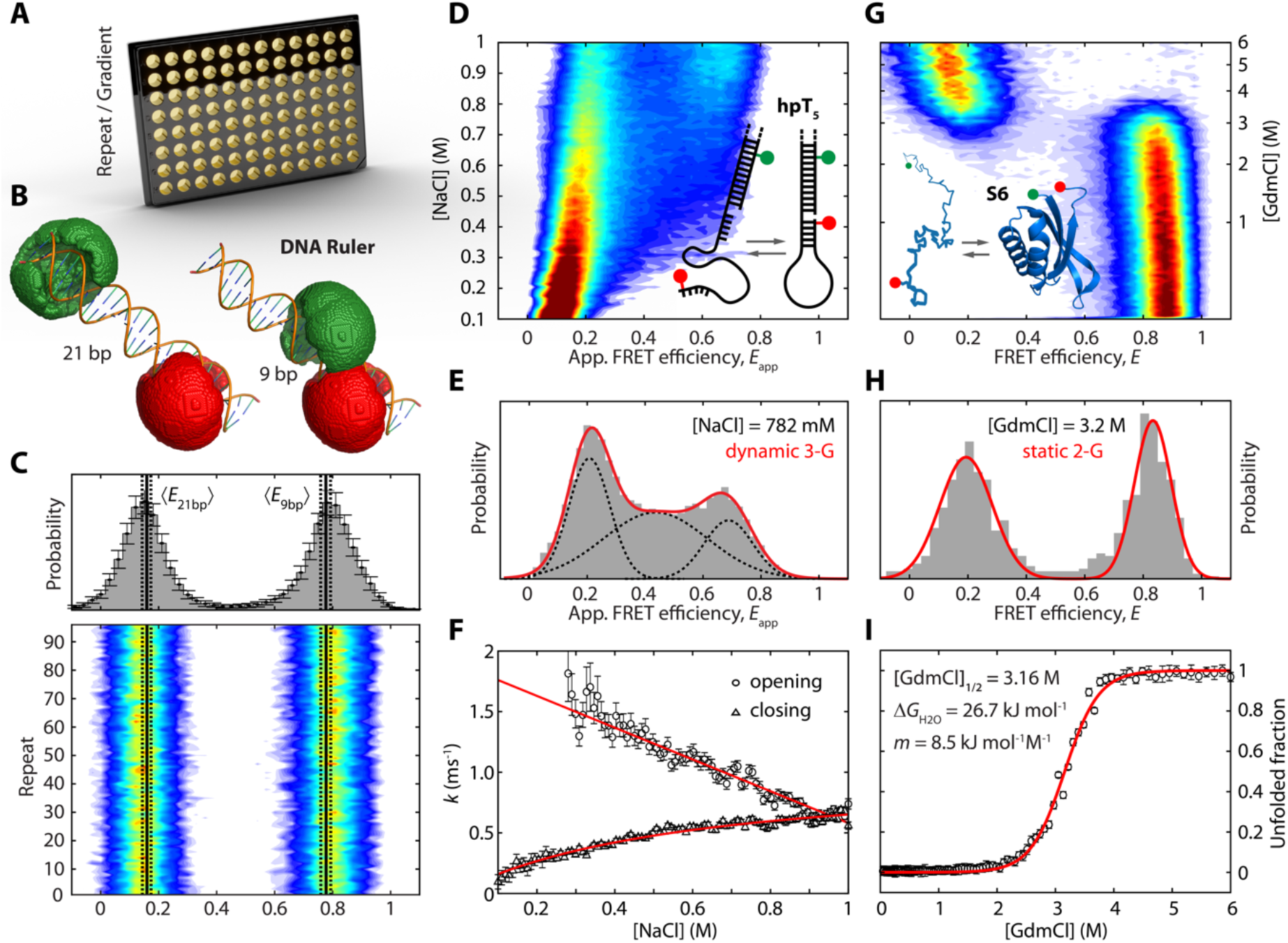
Probing conformational changes of biomolecules by multiwell plate smFRET. **(A)** The multiwell plate format provides the convenience to screen a large number of conditions, such as multiple sample repeats or concentrations gradients. **(B, C)** Evaluation of the accuracy and precision of multiwell plate smFRET measurements using rigid DNA ruler constructs. 96 independent, but identical repeats of a mixture of two double-stranded DNA ruler constructs with 21- and 9-bp spacing between the acceptor (red) and donor (green) fluorophores (see (panel B) were probed in a 96-well plate smFRET measurement (panel C). Shown are cumulated FRET efficiency histograms of all 96 wells (panel C, top) and a two-dimensional (2-D) histogram of FRET efficiency *E* versus multiwell plate repeat (panel C, bottom). The expected FRET efficiencies for the two constructs (*E*_9*bp*_ = 0.777 ± 0.009 and *E*_21*bp*_ = 0.159 ± 0.008), as calculated by accessible volume (AV) simulations^30^, are indicated as black solid lines. The confidence intervals of the AV cloud simulations are shown as black dashed lines. The mean absolute deviation from the theoretical FRET efficiencies of <8.2% for both constructs and the low standard deviation of the mean of *σ*_⟨*E*⟩_ <0.006 demonstrates the high accuracy and precision of the multiwell plate smFRET measurements. **(D)** Salt-dependent structural dynamics of the hpT_5_ DNA hairpin in a 96-well plate. Shown is a 2-D histogram of apparent FRET efficiency (*E*_app_) versus NaCl concentration created from a 96-well plate smFRET measurement. The concentration of salt in the buffer solution was increased in 96 steps from 100 mM to 1000 mM NaCl. A schematic of the hairpin structure is shown, with donor and acceptor fluorophore positions indicated. The hairpin is primarily in the open conformation (*E*_app,O_ ≈ 0.1) at low salt levels. At elevated NaCl concentrations, the closed conformation (*E*_app,C_ ≈ 0.7) becomes increasingly populated. **(E)** FRET efficiency histogram of hpT_5_ at 782 mM NaCl. The intermediate FRET population between the open and closed state originates from molecules changing their conformation between *E*_app,O_ and *E*_app,C_ on the millisecond timescale. The kinetic rates were quantified by fitting a dynamic three-Gaussian (3-G) model (black dashed lines) to the FRET efficiency histogram (red line), as described in refs^31–33^. **(F)** Kinetic analysis of hpT_5_ hairpin dynamics. Opening rates (circles) decrease linearly with increasing NaCl concentration (red straight line). Closing rates (triangles) exhibit a non-linear behavior at lower salt concentrations, which is well described by a model considering the apparent concentration of the complementary strands of the 5 bp proximal stem (red curved line). The error bars indicate the standard deviation of the dynamic 3-G fit. **(G)** GdmCl-induced unfolding of the small globular protein S6 in a 96-well plate format. Shown is a 2-D histogram of FRET efficiency *E* versus GdmCl concentration. The concentration of the denaturant GdmCl in the buffer solution was increased in 96 steps from 0 to 6 M. A schematic of the protein structure is shown, with donor and acceptor fluorophore positions indicated. At [GdmCl] ≈ 3.2 M the protein transitions from a compact folded conformation at *E*_F_ ≈ 0.8 to the unfolded conformation at *E*_U_ ≈ 0.2 following a two-state unfolding mechanism. **(H)** FRET efficiency histogram of S6 at 3.2 M GdmCl (grey bars). The fraction of unfolded molecules was quantified for each GdmCl condition by modeling the corresponding FRET efficiency histogram with a double Gaussian distribution (static 2-G fit, red line). **(I)** Fraction of unfolded S6 molecules as a function of GdmCl concentration. The GdmCl concentration of half occupancy [GdmCl]_1/2_, the transition slope *m*, and the change in free Gibbs energy between the folded and unfolded conformation at zero denaturant Δ*G*_H2O_ = *m* · [GdmCl]_1/2_ were extracted by fitting Eq. 11 (red line) to the data points (circles). The extracted fit parameter values are given in the panel.

### Probing conformational changes of biomolecules by multiwell plate smFRET

Dynamic structural changes of nucleic acids and proteins are central to their functionalities^2^. smFRET is frequently used to gain structural and dynamic insights into conformational changes of nucleic acids and proteins, and to decipher their folding and assembly mechanisms^3,5,8^. In particular changes of solution conditions are an important tool to tune kinetics of structural transitions and to drive molecules into desired conformations. Here, a large dataset is particularly advantageous to extract kinetic and thermodynamic parameters with high accuracy and precision. To demonstrate this, we applied our multiwell plate smFRET assay to a highly dynamic DNA hairpin system and to the unfolding of a small globular protein.

In a first experiment, we determined the effects of salt on the kinetics of a dynamic DNA hairpin (hpT_5_) comprising a 5-bp long complementary annealing stem and a single-stranded loop of 21 thymidine (dT) nucleotides (nts) (Fig. 2D inset, Table S4). To explore the salt-dependent opening and closing rates of hpT_5_, we designed a 96-step gradient of NaCl from 0.1 M to 1 M and added ∼100 pM of the DNA hairpin to each well. At low salt concentrations, the hairpin appeared mostly in the open, low FRET state (*E*_app,O_ ≈ 0.1). Upon increasing salt concentrations, we observed the appearance of a high FRET population (*E*_app,C_ ≈ 0.7) representing the fully annealed DNA hairpin structure. The hairpin was designed such that it is transiently annealed at a temperature of 25 °C even at 1 M NaCl, thereby rapidly interconverting from the closed to the open conformation and back during the ∼1-ms-long passage time through the confocal observation volume (Fig. S5A). Alternating molecular states changing conformation on the millisecond timescale generate an intermediate third FRET efficiency peak, termed bridge population, as illustrated for [NaCl] = 782 mM (Fig. 2E). In order to quantify the interconversion rates, *k*_open_ and *k*_close_, to the open and the closed states of hpT_5_, respectively, we used a dynamic 3-Gaussian (3-G) approximation as described earlier^31–33^. Therefore, we modeled the *E*-histogram for each NaCl concentration with three coupled Gaussian distributions corresponding to the open, bridge-like, and closed population (Fig. 2E, solid and dashed lines, Fig. S5B). The extracted opening and closing rates for the DNA hairpin are plotted in Fig. 2F. We observed a linear salt dependence of the opening rate (red straight line) with a decline of *m*_open_ = −(1.31 ± 0.04) ms^34^M^34^ and an extrapolated rate at zero salt of 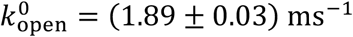

By contrast, the closing rate deviates from a linear behavior and exhibits an unexpected curvature at lower salt concentrations towards smaller rates. To explain this behavior, we developed a model to account for this curvature by considering the apparent local concentration [*A*] of the 5-bp strand around its complementary strand *Ā* (*i*.*e*., proximal stem of hpT_5_ connected by the 21-nt long loop). Briefly, the concentration [A] was calculated from the spherical volume that is spanned by the average distance *R*_A−Ā_ between the two ends of the 21-nt long loop, where *R*_A−Ā_ was derived from the root-mean-square end-to-end distance of a worm-like chain polymer with contour length *l*_c_ = 14.2 nm and a calculated ionic-strength dependent persistence length *l*_*p*_(*I*)^34^ (Eq. 9, *SI*). Strikingly, the salt-dependent increase of the apparent concentration [A], as modelled by this approach, reproduces the observed curvature of the closing rate *k*_close_([NaCl]), as shown in Fig. 2F (curved red line). We find an apparent closing rate of 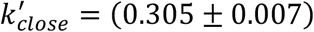· 10^6^ s^−1^M^−1^, which is in good agreement with earlier reports^34^. Furthermore, from the lower and upper limit of the persistence length of the loop (*l*_*p*_(∞) = 0.75 nm and *l*_*p*_(0) = 2.09 nm at infinite and zero concentration of NaCl, respectively) a lower and upper boundary of the hpT_5_ closing rate can be predicted as 0.022 ms^−1^ < *k*_close_ < 1.02 ms^−1^. Taken together, our multiwell plate experiment on the DNA hairpin yielded detailed insights into the molecular kinetics of this dynamic nucleic-acid system, which would be challenging to extract in independent single-well measurements. Especially the curvature in the low-salt regime could be easily missed at reduced sampling density. In fact, we estimate, with bootstrapping, that the variability (*i*.*e*., accuracy) of opening rates decreases with the number of probed conditions. We found that the standard deviation drops from *σ*_5_ = 0.154 ms^−1^ for 5 measured samples to *σ*_70_ = 0.011 ms^−1^ for 70 measured samples, demonstrating that a fine-sampled screen with 70 different conditions decreases the standard deviation by 14-fold.

Besides kinetics, many biophysical studies are interested in thermodynamic stabilities of proteins or protein complexes. One possibility to assess thermodynamic stability of proteins is to titrate the protein with a destabilizing chaotropic reagent or a denaturant in small increments and by fitting the data with a transition function (*e*.*g*., linear extrapolation method, LEM) to extract the denaturant concentration of half occupancy [GdmCl]_1/2_, the transition slope *m*, and the change in free Gibbs energy between the folded and unfolded conformation at zero denaturant concentration Δ*G*_H2O_ = *m* · [*GdmCl*]_1/2_^35–37^. Such measurements benefit from a high sampling density to avoid overparameterization and to increase fitting accuracy and precision.

Here, we demonstrate the power of a dense 96-well plate sampling by equilibrium unfolding of the small globular protein S6 with guanidinium chloride (GdmCl). To this extent, we prepared S6 protein site-specifically labeled with a donor and acceptor fluorophore and subjected the protein to increasing GdmCl concentrations in logarithmic steps from 0 M to 6 M, while maintaining the concentration of S6 at ∼100 pM. We performed multiwell plate smFRET measurements and probed equilibrium unfolding of S6 in a total of 96 steps by probing each condition for 20 min. We plotted the obtained individual *E*-histograms versus GdmCl concentrations in a 2D histogram (Fig. 2G). This denaturation map shows a compact folded conformation of S6 at *E*_F_ ≈ 0.9 at low GdmCl concentrations and the denaturant-induced unfolding of the protein into an expanded conformation at *E*_U_ ≈ 0.2 beyond 2.5 M GdmCl. Unlike in the case of the DNA hairpin, we did not observe an intermediate FRET population, indicating that S6 does not show millisecond transition kinetics, in agreement with earlier reports^38,39^. Fitting each individual FRET efficiency histogram with a double Gaussian function (2-G) (Fig. 2H), we extracted the average FRET efficiencies as well as the fraction of unfolded molecules at increasing GdmCl concentrations (Fig. 2I). We extracted a transition midpoint of [GdmCl]_1/2_ = (3.16 ± 0.01) M and a change in Gibbs free energy of Δ*G*_H2O_ = (26.7 ± 0.7) kJ mol^−1^, in good agreement with earlier reports^39^. Interestingly, the obtained transition slope of *m* = (8.5 ± 0.2) kJ mol^−1^M^−1^ is slightly higher than the reported value of (4.0 ± 0.4) kJ mol^−1^M^−1^, likely originating from the higher pH used in our study (pH 8) as compared to previous studies (pH 6.25)^39,40^. Noteworthy, the high sampling density decreased the standard deviation of the midpoint, *m*-value and thermodynamic stability by 2 to 4-fold (Fig S6D).

In addition to stability, smFRET can also provide structural insights into polypeptide chain properties such as the radius of gyration of the Θ-state (ideal chain), where interactions with the solvent compensate the effect of the excluded volume and the polymer transitions from a globular to a coiled conformation^41^. To demonstrate this on the smFRET data retrieved from S6 unfolding, we extracted the radius of gyration (Fig. S6E) of the unfolded peptide chain of S6 by fitting the Sanchez model to the mean FRET efficiencies of the unfolded state^41^. At a scaling exponent of *ν*= 1/2 (Fig. S6F), we found the radius of gyration of the Θ-state to be *R*_G, Θ_ = (2.38 ± 0.17) nm, remarkably close to the theoretical prediction of 2.21 nm (*SI*). Interestingly, the compaction factor *α* = *R*_G_/*R*_G, Θ_ (Fig. S6G) of our structural analysis reveals an early coil-to-globule transition of S6 at a GdmCl activity below the actual folding transition at *a*_GdmCl_ = 1.73, as also observed for the cold shock protein CspTm and spectrin domain R17^41^.

Taken together, our automated multiwell plate smFRET platform allowed us to explore kinetic and thermodynamic parameters governing biomolecular folding in smFRET experiments at unprecedented resolution. The consistency between the kinetic rates and thermodynamic stability of the DNA hairpin and the protein S6 with previous reports demonstrate the reliability of the multiwell plate approach. Moreover, we have discovered new insights into biomolecular folding mechanisms including the non-linear closing dynamics of the DNA hairpin at low salt concentrations and the early coil-to-globule transition of the protein S6. Given the breadth and depth of information that can be gained from a single multiwell plate smFRET measurement, we anticipate that the acquisition of high-content smFRET datasets using this format will open new possibilities for discovery in biomolecular folding and dynamic structural biology.

### Observing binding modes of multiple proteins to a single substrate by multiwell plate smFRET

The previous examples demonstrated the ability of our platform to precisely sample changes of molecular conformations and kinetics upon altering solution conditions. Another opportunity by a multiwell plate smFRET assay is to explore target binding modes of multiple, competing reaction partners. A prominent example is the competitive DNA binding of the single-stranded DNA binding protein SSB and the DNA strand-exchange protein RecA. Both proteins readily bind to single-stranded DNA (ssDNA), however, their interaction mode is different. SSB occludes 35 or 65-nt-long stretches on ssDNA and dissolves DNA secondary structures^42^. RecA, by contrast, is known to form a filament on ssDNA with a 3-nt footprint, and is a key player in homologous recombination.^43^ Single-molecule experiments have discovered already a direct interaction and competition of RecA and SSB, with RecA nucleation being facilitated by SSB, likely by RecA–SSB complexes^44–46^. However, this intricate interactive behavior and likely multiple pathways of binding make it difficult to explore the full parameter space of affinities and nucleation by sets of single-well measurements. The multiwell plate format offers the possibility to apply concentration gradients of two molecules against each other (Fig. 3A), making it easier to identify competitive and cooperative effects as well as to pinpoint a reaction scheme and to extract dissociation constants. To demonstrate this capability, we performed multiwell plate smFRET measurements of SSB and RecA and studied interactive binding of both proteins to ssDNA.

**Figure 3.**
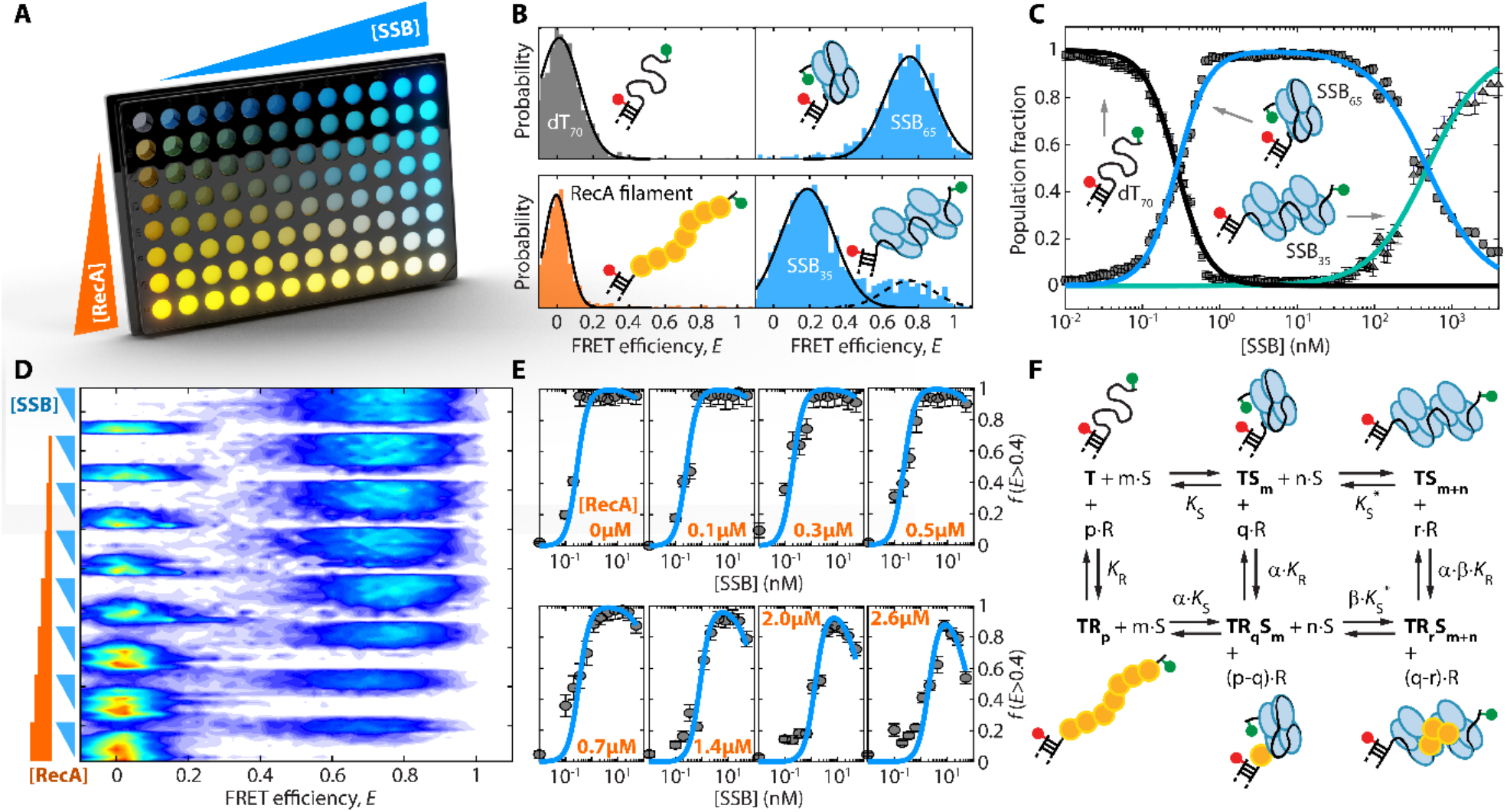
Observing binding modes of multiple proteins to a single substrate by multiwell plate smFRET. **(A)** Competitive binding of RecA and SSB to ssDNA probed in a 96-well plate measurement by a combined variation of the concentration of SSB (0 to 50 nM) and RecA (0 to 2.6 μM) using the DNA construct dT_70_ as a substrate. **(B)** The DNA construct dT_70_ consists of 18-bp double-stranded DNA region carrying an acceptor fluorophore and a 70-nt long thymidine (dT_70_) ssDNA overhang labeled at the terminal end with a donor fluorophore (top left panel). This design allows distinction of two major binding modes of the SSB tetramer, SSB_65_ and SSB_35_ (top and bottom right panel) with FRET efficiencies of *E*_65_ ≈ 0.8 and *E*_35_ ≈ 0.2, respectively, as well as the RecA filament (bottom left panel) with a FRET efficiency of *E*_RecA_ ≈ 0. The dT_70_ construct in the absence of any protein exhibits a broad FRET efficiency distribution at *E*_dT70_ ≈ 0.05. **(C)** Speciation curves as obtained from a multiwell plate smFRET experiment of dT_70_ subjected to increasing concentration of SSB ranging from 0 to 4 μM (in the absence of RecA). The resulting population fractions, extracted from FRET efficiency histograms, enabled calculation of the SSB concentrations at half occupation and the respective Hill coefficients (see main text). **(D)** 2D FRET efficiency histogram of the competitive binding of RecA and SSB to dT_70_ as obtained from a 96-well plate smFRET measurement. Bars and wedges on the left side depict the RecA and SSB concentrations. In the absence of RecA (top rows), a transition from the dT_70_ (*E*_dT70_ ≈ 0.05) state to the SSB_65_ state (*E*_65_ ≈ 0.8) can be observed at low SSB concentrations. With increasing RecA concentration this transition shifts to higher concentrations of SSB, demonstrating that RecA inhibits SSB from binding to dT_70_. Interestingly, at the two highest RecA concentrations, the high FRET population shifts to lower FRET values, indicating a more extended conformation of dT_70_, most likely originating from a mixed binding of RecA and SSB. **(E)** Fraction of molecules with FRET efficiency *E* > 0.4 versus [SSB] for increasing RecA concentrations. The sigmoidal curves show three different correlations with [RecA]: i) *c*_1/2_(SSB) shifts to higher values; ii) *f*_max_ decreases, and iii) the fraction at maximal SSB concentration declines. **(F)** Model of interconverting states used to describe competitive binding of SSB and RecA to dT_70_. T: empty dT_70_; TS_m_: SSB on dT_70_ in SSB_65_ binding mode; TS_m+n_: SSB on dT_70_ in SSB_35_ binding mode; TR_p_: RecA filament on dT_70_; TR_q_S_m_: mixed state of SSB_65_ and RecA on dT_70_; and TR_r_S_m+n_: mixed state of SSB_35_ and RecA on dT_70_. The values *K*_*S*_, 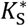, *K*_R_, *α* · *K*_R_, *α* · *β* · *K*_R_, *α* · *K*_*S*_, *β* · 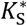 denote the apparent dissociation constants with the corresponding Hill coefficients *m, n, p, q*, and *r*. The RecA concentrations at half occupation 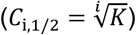 was extracted from the fit in panel E (blue curves). Values are given in the main text.

We designed a DNA construct with an 18-bp double-stranded DNA region carrying an acceptor fluorophore and a 70-nt long thymidine (dT_70_) ssDNA overhang on the 3’-end terminated by a donor fluorophore (Fig. 3B inset, Table S4). The proximity of the donor and acceptor fluorophores allows monitoring the compaction of the dT_70_ overhang upon binding of SSB or RecA (Fig 3B). In absence of any protein, the ssDNA exhibits a low FRET efficiency (*E*_dT70_ ≈ 0.05) as the DNA is only weakly collapsed. In the presence of low concentrations of SSB (*e*.*g*., 1 nM), SSB binds to the ssDNA in the SSB_65_ binding mode, where 65 nts are occluded by SSB, leading to a FRET efficiency of *E*_65_ ≈ 0.8 (Fig. 3B). With increasing SSB concentrations (> 50 nM), the SSB_65_ binding mode transitions into the SSB_35_ binding mode, incorporating two tetramers of SSB (Fig. 3B) and, hence, leading to a more expanded dT_70_ conformation with a FRET efficiency *E*_35_ ≈ 0.2. RecA, in presence of ATP, forms a nucleoprotein filament, thereby stretching ssDNA. Hence, binding of RecA to dT_70_ results in its elongation beyond the dynamic range of FRET, yielding a narrow distribution of FRET efficiencies of *E*_RecA_ ≈ 0. Notably, the signature of RecA binding is clearly distinguishable from the two major binding modes of SSB (Fig. 3B)^47^. In summary, the high FRET contrast of the three states, SSB_65_, SSB_35_, and RecA filament, allows shedding light on the interactive behavior of both proteins in the presence of ssDNA.

In a first experiment, we studied SSB binding to ssDNA alone. To this end, we performed a multiwell plate measurement of dT_70_ (∼100 pM) subjected to increasing SSB concentration ranging from 0 to 4 μM (Fig S7). We determined the fractions of molecules in the dT_70_ (rectangle), SSB_65_ (circle) and SSB_35_ (triangle) state as a function of SSB concentration (Fig. 3C). The fractions were then modeled by Eq. 15 (*SI*) to derive the concentrations at half occupancy of *c*_*S*,1/2_ = (278 ± 1) pM and 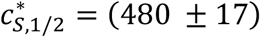 nM and the corresponding Hill coefficients of *m* = (2.12 ± 0.07) and *n* = (1.19 ± 0.04) for the SSB_65_ and SSB_35_ binding mode, respectively. The rapid continuous mapping across different concentrations agrees well with the simple binding theory (Eq. 15) and earlier reports^42,44,48^.

Subsequently, we performed a multi-well plate measurement of dT_70_ with 96 different combinations of [RecA] and [SSB] to study the competition between RecA filament formation and SSB_65_ binding. To this end, we varied the SSB concentration in 12 steps from 0 to 50 nM along the columns of the plate and the RecA concentration in 8 steps from 0 to 2.6 μM along the rows of the plate (Fig. 3A). In the measurements without RecA (top row, Fig. 3D), the transition from the broad dT_70_ state (*E*_dT70_ ≈ 0.05) to the SSB_65_ binding mode (*E*_65_ ≈ 0.8) appears at a low concentration between 0.2 and 0.35 nM agreeing well with our previously determined *c*_*S*,1/2_ = 0.28 nM, and the independent observation of the change in fluorescence anisotropy of the acceptor (Fig. S8C). With increasing RecA concentrations, the occupation of SSB_65_ shifted to higher SSB concentrations. At the same time, the RecA population at *E*_RecA_ ≈ 0 became more abundant, reflecting a modulation of the apparent dissociation constant of SSB by competitive binding of RecA. Interestingly, at high RecA concentrations ([RecA] = 2 − 2.6 μM) a shift of the high FRET population to lower FRET efficiencies is observed, which suggests a new state of combined RecA and SSB binding.

For a quantitative analysis of the interactive binding of SSB and RecA to ssDNA, we extracted the fractions of DNA molecules bound in the SSB_65_ mode *f*(*E* > 0.4) from the 2D histogram (Fig. 3E). As qualitatively observed, at higher RecA concentrations, the transition to the SSB binding mode occurred at higher SSB concentrations. Surprisingly, we observed a drop of the maximal fraction of SSB_65_ at [RecA] > 0.5 μM and [SSB] > 10 nM. The depopulation of the SSB_65_ state at elevated [RecA] supports the presence of mixed RecA–SSB states. Considering the two SSB binding modes and the presence of a RecA-filament, we build a 6-state model (Fig. 3F), which contains the known dT_70_ (T), RecA filament (TR_p_), SSB_65_ (TS_m_) and SSB_35_ (TS_m+n_) states, as well as the two mixed states of RecA-SSB (TR_q_S_m_) and RecA-2xSSB (TR_r_S_m+n_). Here, the values *K*_*S*_, 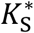, *K*_R_, *α* · *K*_R_, *α* · *β* · *K*_R_, *α* · *K*_*S*_, *β* · 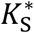 denote the respective dissociation constants with the corresponding Hill coefficients *m, n, p, q*, and *r*. The 6-state reaction scheme allowed us to model the fractions of molecules with *E* > 0.4 (Fig. 3E). Since it was unclear where the states TR_q_S_m_ and TR_r_S_m+n_ appear on the FRET axis, and thus what states contributed to *f*(*E* > 0.4), we performed 75 different fittings with varying state and probability configurations using

Eq. 16 (*SI*) and the previously determined values 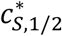, *m* and *n*. For each combination of state and probability configurations, the reduced chi-squared 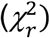 was calculated from the residual, taking the number of fitting parameters into account. We found the smallest 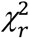 value 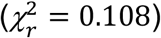 for the full reaction scheme involving all 6 states, where *f*(*E* > 0.4) describes the combined fraction of state TS_m_ (SSB_65_) and TR_q_S_m_ (RecA-SSB) (Fig. S10). Looking at the formation of a RecA-filament, we find half concentrations of occupancy of *c*_R*b*,1/2_ = (425 ± 91) nM, and for RecA-SSB formation a value of *c*_R*q*,1/2_ = (237 ± 76) nM, and for RecA-2xSSB a value of *c*_R*r*,1/2_ = (278 ± 94) nM with the corresponding Hill coefficients of *p* = (4.9 ± 1.5), *q* = (1.9 ± 1.5) and *r* = (3.1 ± 1.6). Hence, our data revealed that RecA affinity is increased 1.8-fold by the presence of a SSB_65_-complexed ssDNA and 1.5-fold by a SSB_35_-complexed DNA. SSB facilitating RecA filament formation was previously observed^44–46^, yet, it was impossible to quantify this enhancement. Taken together, our measurements illustrate the possibility to screen three or more component systems for cooperativity or competition within a single 96-well plate smFRET experiment and, by extension, reveal new, unexpected cooperativities and competitions.

### Multiwell plate smFRET screening of drug–protein interactions

smFRET experiments are increasingly employed to study the molecular mechanisms of small-molecule binding to target proteins in a variety of applications, ranging from enzyme–ligand interactions to probing reversal effect of small molecule corrector compounds on protein misfolding^49–53^. However, larger-scale screenings of molecular compounds by smFRET, as used in pharmacological research and drug discovery, have been limited because tools to conduct such time- and labor-intensive measurements are lacking. For example, recently, we used smFRET to study the misfolding and drug rescue mechanism of the cystic fibrosis transmembrane conductance regulator (CFTR), an ion channel protein that is defective in people with cystic fibrosis (pwCF). We used a minimal hairpin model derived from the CFTR transmembrane helices 3 and 4 (TM3/4) carrying a patient-derived mutation and found that misfolding induced by the point-mutation V232D in TM3/4 could be rescued with the drug Lumacaftor^54^. Such experiments required tens of single chamber smFRET measurements with extensive cleaning steps, long equilibration periods, and repeated sample reconstitution for single conditions. Extension of such measurements to larger-scale screenings and would benefit massively from an automated multiwell plate format in order to probe multiple small molecules or multiple patient-derived mutations.

Here, we explored such an automation for molecular screening of drug–protein interactions by our multiwell plate smFRET assay. The screen comprised four protein variants and two small molecule compounds. The protein variants consisted of wildtype TM3/4 (WT) and three mutant variants E217G, Q220R, and V232D TM3/4 (Fig. 4A, B and Tab. S4). All three mutations are CF-phenotypic and known to cause maturation defects and misfolding of CFTR^55,56^. The drug molecules were two CFTR interacting correctors, Lumacaftor (VX-809) and Galicaftor (ABBV-2222)^57^. Lumacaftor as well as Galicaftor have been developed to rescue the most common misfolding mutation ΔF508 in CFTR^58,59^, but also showed improved maturation of CFTR with V232D^55,60^. To read out misfolding and drug rescue of the transmembrane helices, we attached donor and acceptor fluorophores close to the N- and C-termini of our TM3/4 hairpins and reconstituted the TM3/4 hairpins in lipid vesicles (Fig. 4B). After reconstitution and transfer into a 96-well plate, we read out the degree of misfolding and partial insertion by collecting FRET efficiency histograms in presence of increasing concentrations of corrector compounds (over each row of the 96-well plate). In this context, a high FRET efficiency is related to correct insertion, while a lower FRET efficiency indicates partial insertion and misfolding^54^. Hence, we can detect misfolding, and by performing concentration screenings of corrector compounds, we can detect the degree of drug rescue of the hairpin structures by reading out FRET efficiencies and determine an *EC*_C5_ of the drug–protein interaction from dose–response curves.

**Figure 4.**
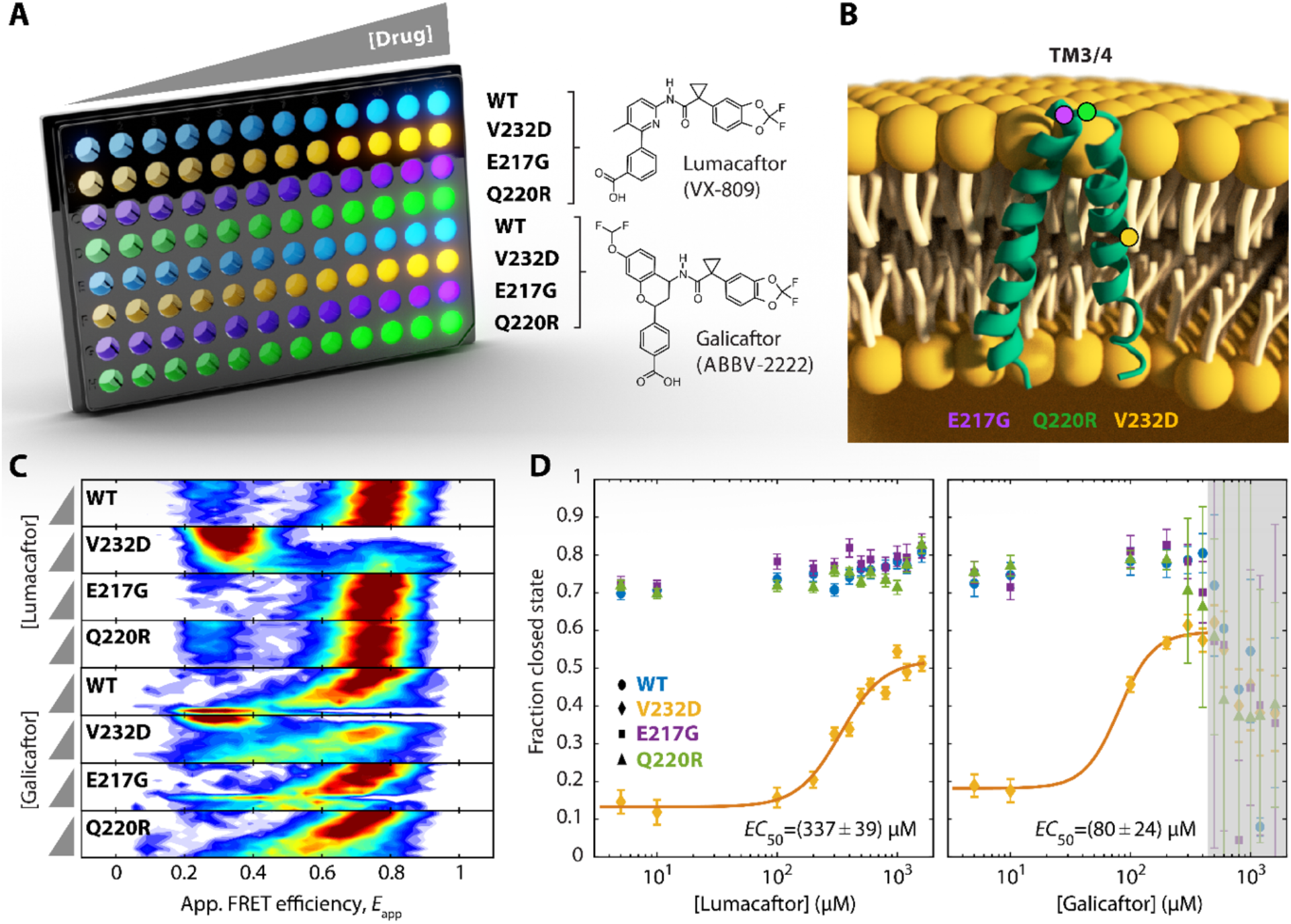
Multiwell plate smFRET screening of drug–protein interactions. **(A)** Titration of CFTR TM3/4 hairpins (WT, V232D, E217G, Q220R) with Lumacaftor (VX-809) and Galicaftor (ABBV-2222) over 12 different drug concentrations in a 96-well plate. To probe TM3/4–drug interactions by multiwell plate smFRET, hairpins were reconstituted in lipid vesicles. **(B)** Schematic of the TM3/4 hairpin structure in a lipid bilayer. The positions of the transmembrane mutant V232D (orange) and of the two loop mutations E217G (purple) and Q220R (green) are indicated by colored circles. **(C)** 2-D histogram of FRET efficiency versus drug concentration for WT, V232D, E217G and Q220R TM3/4 in the presence of either Lumacaftor or Galicaftor. The concentration of the drugs was increased in 12 steps from 5 to 1600 μM (see grey wedges). The misfolded mutant V232D TM3/4 exhibits folding recovery in the presence of both drugs as seen by a shift from a low FRET to a high FRET population (*i*.*e*., transition from an open to a closed hairpin conformation). The other TM3/4 variants remain in a high FRET state (closed conformation) over the entire concentration range, indicating no effect on the hairpin conformation. The shift and extra broadening of the FRET population at high concentrations of Galicaftor originates from an increase in fluorescence background. **(D)** Dose–response curves for TM3/4–drug interactions. The plots depict the fraction of TM3/4 molecules in the closed conformation versus Lumacaftor (left panel) or Galicaftor concentration (right panel) for WT, V232D, E217G and Q220R. The fractions were extracted by Gaussian fitting of the FRET efficiency histograms (Fig. S11) shown in panel C. To determine the effective concentration of half occupancy for the V232D TM3/4 mutant, the dose-response curves were fitted with a Hill-type function to yield *EC*_50_ values of *EC*_50_ = (337 ± 39) μM and *EC*_50_ = (80 ± 24) μM for Lumacaftor and Galicaftor, respectively. In the case of Galicaftor, data points above 500 μM were excluded from the fit, due to deviation of the fluorescence lifetime of the donor and acceptor (indicated in grey).

Using our multiwell plate assay, we collected FRET efficiency histograms for our four hairpin variants and two corrector molecules (Fig. 4C and S11). The histogram for WT TM3/4 showed, in the absence of any corrector, a compact structure with a high fraction of molecules being in the correctly folded, high FRET efficiency state (Fig. 4D, blue dots). Upon addition of either Lumacaftor or Galicaftor, little change of the degree of folding was observed. Also the variants E217G and Q220R TM3/4 mostly retained a closed conformation with high FRET efficiency similar to WT TM3/4, in agreement with our earlier study^61^. V232D TM3/4, on the other hand, in the absence of any corrector, appeared misfolded and adopted a mostly open conformation with low FRET efficiency (Fig. 4C, D). Titration of Lumacaftor restored folding of V232D TM3/4 to a large degree with *EC*_50_ = (337 ± 39) μM, which is in excellent agreement to our previously reported *EC*_50_ of 347 μM (Fig. 4D)^54^. Lumacaftor, however, showed no drastic effect on the loop mutants E217G and Q220R TM3/4 as they remained largely folded even in the absence of the corrector, and we observed only a slight stabilization at high concentrations. In the case of Galicaftor, which we did not study previously, we observed that V232D TM3/4 could be rescued from misfolding as well, while the other variants were little affected by increasing Galicaftor concentrations (Fig. 4C, D). Noteworthy, at [Galicaftor] > 400 μM, we detected a deviation of the fluorescence lifetime of the donor and acceptor, which interfered with the detection and, thus, was not considered in our analysis (Fig. 4D, grey area). Interestingly, the observed *EC*_50_ for the rescue of V232D TM3/4 was *EC*_50_ = (80 ± 24) μM and, thus, much lower than for Lumacaftor. Recent *in vivo* experiments also observed that Galicaftor is 12-fold more potent than Lumacaftor in rescuing full-length V232D CFTR at the plasma membrane^60^. This corroborates that the structural readout of smFRET experiments on TM3/4 misfolding and its rescue by small molecule correctors provides insights into drug-action mechanisms.

In summary, our experiments here on TM3/4 hairpin screening illustrates that within a single, automated multiwell plate smFRET experiment, we were able to recover structural information of misfolding events and their rescue, illustrating the suitability of our assay for drug screenings. We anticipate that such a multiwell plate assay opens up new avenues to use smFRET for a characterization of patient–derived mutations on conformational dynamics and their rescue, thus new approaches for rational drug design and drug discovery.

## Conclusions

Here, we introduced a platform for automated smFRET experiments in a multiwell plate format. With different examples, we illustrated how accurate and precise FRET efficiencies as well as conformational dynamics, molecular competitions, and information on small–molecule–protein interactions can be obtained from multiwell plate measurements. Along with a detailed description of the hardware components, which are all commercially available and can be installed without expert knowledge, we provide an open-source software suite for data acquisition, analysis, and visualization (see *SI* for details), offering an easily adaptable approach for other labs to integrate the multiwell plate smFRET assay into their workflows.

All our examples were performed on 96-well plates with a 20-min measurement time per well. This accumulates to a total measurement time of 32 h and was sufficient to collect extensive data for all provided examples. However, our assay can be easily adjusted to either smaller (*e*.*g*., 48-wells) or larger (384-wells) multiwell plate formats, as desirable for the application. Further, the data acquisition software allows to select the specific wells to be probed on the multiwell plate, such that only a subset of conditions can be probed, and in the case of varying measurement statistics, the data acquisition time can be adjusted flexibly. With respect to measurement time, we noted that 32 h measurements can cause loss of fluorescent molecules, *e*.*g*. by non-specific adsorption to the 96-well plate. However, such loss can be prevented by the addition of a small percentage of surfactant (*e*.*g*., Tween20) or a brief preincubation with BSA to achieve surface passivation.

One important aspect for multiwell plate measurements is a swift sample preparation. While all of the described examples were manually pipetted, in an optimized pipetting scheme taking approximately 2 h to prepare an entire plate, commercial implementations of micro-dispensers allow to fill a 96-well plate within 15 min in an automated fashion (*SI*). Such a rapid, reliable preparation of 96-well plates provides an important step towards smFRET high-throughput screenings. In fact, the ease of setting up such multiwell plate experiments will unleash the unique possibility to extensively, yet swiftly, bridge the gap between structural and functional aspects of biomolecular systems in dynamic structural biology and biophysics, and beyond.

Overall, with our automated multiwell plate platform, we open up unique possibilities to use high-content smFRET data for biomolecular screening and drug discovery. The increased sampling of multiwell plate smFRET, as demonstrated in our work, allows, for instance, to (i) discover unexpected interactions in multi-component systems; (ii) screen many different small molecules for affinities and effects on molecular conformations; and (iii) discover subtle conformational changes, which are typically inaccessible in single-well measurements due to low parameter space sampling. Together with our open-source cross-platform software suite and the easy-to-implement additions to already available smFRET setups, we anticipate that multiwell plate smFRET will enable the community to set up such a system in their labs and to gain in-depth insights into biological systems, spanning from protein folding to nucleic-acid structures to protein-drug interactions.

## Material and Methods

### DNA and protein preparation

DNA samples, including fluorescently labeled oligos, were commercially obtained and annealed according to the details given in the *SI*. Proteins were either purchased commercially or expressed recombinantly and purified. Details are given in the *SI*.

### DNA ruler experiments

Annealed DNA rulers were diluted to ∼100 pM in buffer (20 mM Tris-HCl, pH 8, 50 mM NaCl) and distributed into each well (Tab. S5).

### DNA hairpin experiments

Annealed DNA hairpin hpT_5_ was diluted to ∼100 pM in buffer (20 mM Tris-HCl, pH 8) with well-specific NaCl concentrations ranging from 100 to 1000 mM in each well (Tab. S5 and S7).

### S6 unfolding experiments

S6 was diluted to ∼100 pM in buffer (50 mM Tris-HCl pH 8, 150 mM NaCl, 2 mM TCEP) with well-specific GdmCl concentration ranging from 0 to 6 M (Tab. S5 and S8).

### RecA and SSB experiments

DNA dT_70_ was diluted to ∼100 pM in buffer (50 mM Tris-acetate pH 7.7, 5 mM Mg-acetate, 50 mM Na-acetate) and SSB (Promega Corporation, USA) was supplemented in specified concentrations (Tab. S5 and S9). For the SSB–RecA competition, the buffer was supplemented with 16 mM ATP and RecA (New England Biolabs, USA) as well as SSB gradients were added to the respective rows (Tab. S10).

### TM3/4 experiments

Reconstituted TM3/4 variants in POPC LUVs were diluted to ∼100 pM in buffer (50 mM Tris-HCl pH 7.4) and added to each well. Wells were supplied with a gradient of small molecule concentrations ranging from 5 to 1600 μM (Tab. S5 and S11).

### Automated multiparameter single-molecule detection setup

Experiments were carried out using a single-molecule confocal fluorescence microscope with pulsed-interleaved excitation and fluorescence anisotropy detection as shown in Fig. 1 and described in detail in the *SI*. Briefly, the microscope was equipped with a motorized sample stage (ASR100B120B, Zaber Technologies, Canada), a heating pad (Lerway, China), and an autofocus system (Perfect Focus System, Nikon, Japan). The immersion water was supplied by a liquid dispenser (Märzhäuser Wetzlar, Germany). The multiwell plates used in this study were glass bottom 96-well plates from IBL Baustoff+Labor GmbH, Austria.

### smFRET data analysis

Data analysis was performed using pyBAT and pyVIZ as well as customized scripts for quantification of center positions, fractions, kinetic rates, and other extracted parameters. Single-molecule events were identified from the acquired photon stream by a burst search algorithm. Details about the primary and secondary data analysis procedures are given in the *SI*.

### Software

The software used in this manuscript is freely available from the GitHub repository: https://github.com/SchlierfLAB/autoFRET. The Python-based platform software suite autoFRET comprises three components: Data Acquisition (pyMULTI), Analysis (pyBAT) and Visualization (pyVIZ). A detailed description of the software package can be found in the SI and on GitHub, including an example data set.

## Acknowledgements

We thank all members of the Schlierf lab for lively discussions during the development of this project. This research was funded by TU Dresden core funds, the DFG (SCHL1896/3-1 and SCHL1896/4-1), the BMBF (OptiZeD Grant Z22E511 (to MS)), the European Social Fund (ESF) and co-financed by tax funds based on the budget approved by the members of the Saxon State Parliament (to MSche) and by a grant from Mukoviszidose Institut gGmbH, Bonn, the research and development arm of the German Cystic Fibrosis Association Mukoviszidose e.V. (to MS). GK acknowledges support the European Research Council (ERC) under the European Union’s Horizon 2020 Framework Programme through the Marie Skłodowska-Curie Grant MicroSPARK (agreement no. 841466) (GK), the Herchel Smith Funds of the University of Cambridge, and the Wolfson College Junior Research Fellowship.

## Competing interests

The authors declare no competing interests.

### Author contributions (CRediT scheme)

|  |  |
| --- | --- |
| Conceptualization | MS; AH; GK; MSche |
| Methodology | AH; KS; PS |
| Software | PS; AH; KS |
| Validation | KS; AH; PS |
| Formal analysis | AH; KS |
| Investigation | KS; NC; MSche; AH |
| Resources | GK; AH; MSche; |
| Data Curation | AH; KS; MSche; NC |
| Writing - Original Draft | AH; NC; MS; MSche |
| Writing - Review & Editing | MS; AH; NC; MSche; KS; GK; PS |
| Visualization | AH, MSche |
| Supervision | MS; AH |
| Project administration | AH; MS |
| Funding acquisition | MS; MSche; GK |

