## Supporting Information for "Farewell to single-well: An automated single-molecule FRET platform for high-content, multiwell plate screening of biomolecular conformations and dynamics"

**This PDF file includes:**

Supporting Methods
Figures S1 to S11
Tables S1 to S11
SI References

### Supporting Methods

#### Automated multiwell plate, multi-parameter fluorescence detection setup for 96-well plate measurements

Figure S1 shows a detailed illustration of the multiwell plate, multi-parameter fluorescence detection (MFD) single molecule detection setup, including the controller and data acquisition hardware. All optical parts were assembled on an optical table (M-VIS3660-SG4-325A, Newport, Irvine, USA). For donor and acceptor excitation, we used two pulsed laser sources with wavelengths of 530 nm (LDH-P-FA-530L, PicoQuant, Berlin, Germany) and 640 nm (LDH-D-C-640, PicoQuant) driven in pulsed interleaved excitation (PIE) mode at a repetition rate of 50 MHz frequently changing between direct excitation of the donor and acceptor fluorophore. In order to obtain a $TEM_{00}$ mode of the laser profile, the linear polarized laser beams were combined into a polarization-maintaining single-mode optical fiber (P3-488PM-FC-2, Thorlabs, Newton, NJ, USA). The resulting beam was collimated (60FC-T-4-RGBV42-47, Schäfter und Kirchhoff, Hamburg, Germany) at the end of the patch cable and directed into the inverted Nikon microscope (Nikon Eclipse Ti, Nikon). We used an average laser power of 110 µW for the green and 90 µW for the red laser, measured before the objective. The fluorescence emission of the donor and acceptor was collected by a 60x water objective (Fl Plan Apo WI 60x with NA 1.2, Nikon, Tokyo, Japan) 30 µm in solution, separated from excitation light by a dichroic mirror (zt532/642rpc, Chroma, Bellow Falls, USA), and focused on a 50 µm pinhole. The confocal volume emission was then separated according to the light polarization by a polarizing beam splitter (PBS: CM1-PBS251, Thorlabs, Newton, USA). Subsequently, in each polarization arm (parallel || and perpendicular ⊥), the donor and acceptor emission were sorted using a dichroic mirror (FF650-Di01, Semrock, USA). Finally, the band-pass filtered (F1: FF01-582/75, Semrock; F2: ET700/75M, Chroma) light was collected by four single-photon avalanche diodes (APD: SPCM-AQR, Excelitas, Waltham, MS, USA) and recorded by four independent channels of a TCSPC module (HydraHarp 400, PicoQuant). A detailed description of a typical MFD setup can be also found in ref [1].

The 96-well plate reading functionality was realized by a motorized x-y stage (Table S1, Zaber Technologies Inc., Vancouver, BC, Canada) mounted onto the microscope body. In order to prevent evaporation and condensation during the measurement time, the 96-well plate was sealed with a parafilm and a heating pad (Table S1), which was operated a few degrees Kelvin above room temperature. The focus was maintained 30 µm in solution using a commercial autofocus system (Perfect Focus System, TI-ND6-PFS, Nikon). The immersion water of the objective was frequently replaced by a liquid dispenser system (Table S1, Märzhäuser Wetzlar GmbH & Co. KG, Wetzlar, Germany). The objective adapter for the dispenser was customized for our water objective by Märzhäuser.

**Open-source software package autoFRET for hardware control and data analysis**

The autoFRET Python software package consists of three individual applications for hardware control of single- or multiwell measurements (pyMULTI), single-molecule data analysis of raw data (pyBAT_otf/pyBAT), and inspection of analysis results (pyVIZ). The software suite is available on GitHub: https://github.com/SchlierfLAB/autoFRET . A detailed description including a link to sample data is given on GitHub. For each program a graphical user interface (GUI) is provided (Fig. S1).

In the measurement GUI pyMULTI, the user can select measurement wells, can choose the acquisition, and can save a project description. The purpose of the script is to automatically control the motorized scanning stage (ASR100B120B-E03T3A), to communicate with the TCSPC module (HydraHarp 400) and to save raw data in an organized folder structure. During a multiwell measurement, for each well a separate folder (*e.g.*, “A01”) is created containing one-minute long measurement files (currently in the HT3 format). These files have a header free binary format (PicoQuant) with 32-bit entries, where each entry encodes the arrival time (macrotime), fluorescence delay time (microtime, 16 ps resolution), and channel (detector 1 to 4) of a single detected photon. The script can be readily adjusted to different hardware if DLLs are provided.

While the analysis software pyBAT_otf (on-the-fly) runs parallel to an ongoing multiwell measurement, pyBAT can be performed after a measurement session as well. Both BAT versions identify fluorescence bursts in the recorded photon streams and thereby extract MFD parameters of single molecules. In the GUI, several settings like microtime gates, inter-photon time thresholds and minimum number of photons per burst can be made according to the repetition rate of the lasers and count rate of the fluorophores. The extracted MFD parameters are stored with a burst identification number in new binary files in the corresponding folder. For larger measurements, we highly recommend the multi-processing option of pyBAT, which improves the runtime (Tab. S2).

The pyVIZ GUI provides the user with multiple visualization options of the extracted MFD parameters (*i.e.*, 2D histograms for multiwell overviews), individual histograms, and scatter plots for cross correlation of two parameters. Additionally, a vast number of filter options like ALEX-2CDE and FRET-2CDE facilitates data interpretation, like *e.g.*, the extraction of double-labeled molecules and the identification of conformational dynamics. Furthermore, the application holds the opportunity to export filtered data in CSV-files and plots in SVG and PNG format.

The software package autoFRET was tested on macOS Monterey and Windows 10. The analysis software (pyBAT/pyBAT_otf) requires 8 GB of memory and benefits from modern CPUs. It will run on older systems as well, but the runtime can drastically increase depending on the processor. Table S2 summarizes the runtimes of pyBAT on several machines.

We provide YML-files to install the required external libraries in an anaconda (https://anaconda.org) environment. The installation can be run manually or by running the provided autoinstall.py script. More advanced users can also install all required libraries (Tab. S3) manually. After all libraries are successfully installed, execute one of the main program files (pyMULTI.py, pyBat.py/pyBat_otf.py, pyVIZ.py). Additional information can be found in the corresponding GitHub Wiki page.

**Oligonucleotide design and labeling**

Fluorescently labeled and HPLC-purified DNA oligos were obtained from IBA (Göttingen, Germany) modified with donor and acceptor fluorophore derivatives of ATTO 532 and ATTO 647N (ATTO-TEC, Siegen, Germany). For the DNA hairpin hp$T_5$, the top strand was terminal labeled at an amino-C6-modified 5'-phosphate-thymidine and the bottom strand was base-labeled at position 25 to an amino-C6-modified thymine (Table S4). In the case of the DNA ruler constructs, the top and bottom strands were base-labeled, while for the d$T_{70}$ construct, both strands were terminal labeled (Table S4). Prior to a multiwell plate experiment, complementary DNA constructs were annealed at a concentration of 5 µM in 10 mM Tris-HCl pH 8.0 (Roth,

Karlsruhe, Germany), 50 mM NaCl (Roth) and 1 mM EDTA (AMRESCO, Solon, OH, USA) by heating to 90 °C and cooling to room temperature in 1 K/min steps.

**Protein design, purification and labeling**

A double-Cys variant of the 101-amino-acid-residue protein S6 from *Thermus thermophilus* was constructed (Tab. S4), as described previously[2]. The protein was site-specifically labelled via thiol–maleimide chemistry with maleimide-functionalized donor (ATTO 532) and acceptor (Abberior STAR 635P, Abberior GmbH) fluorophores, following standard procedures. The labelled protein was separated from unbound fluorophores by size-exclusion chromatography.

Protein expression and purification of CFTR TM3/4 hairpins were performed according to our previous protocols with minor adjustments[3]. To introduce point mutations, the sequence of the TM3/4 fusion protein construct was changed by site-directed mutagenesis (QuikChange Lightning Multi kit, Agilent, USA), where a C-terminal His-tag was deleted, and an additional enterokinase cleavage site was introduced (Fig. S3). Expression in *E. coli* BL21(DE3) and in M9 minimal medium was executed as described earlier[3], the purification protocol was modified as follows: After cell lysis and IMAC, the elution fractions were concentrated with Amicon™ Ultra-15 centrifugal filter units (10 kDa molecular weight cutoff, Merck, Germany) at 5000 x *g*, 10 min, 4°C before heat treatment, buffer exchange into labeling buffer (50 mM Tris, 150 mM NaCl, 0.1% (w/v) Triton X-100, 1.5 mM TCEP, pH 7.25) and thrombin digestion. Before fluorescent labeling, reverse IMAC was performed with His SpinTrap columns (Cytiva, USA) following the manufacturer's gravity protocol: Labeling buffer supplemented with 5 mM imidazole was used for column equilibration before application of 600 µL of digested sample, supplemented with 5 mM imidazole, to the column. The sample was incubated with the column for 15 min at room temperature and then recovered by centrifugation at 100 x g for 30 s. The flow-through with cleaved TM3/4 was collected and analyzed with SDS-PAGE. Labeling of cleaved TM3/4 variants was carried out with maleimide-functionalized dyes ATTO 532 (donor) and ATTO 647N (acceptor) with 200-fold molar excess of donor dye was added to the sample and incubated for 15 min at room temperature before adding 50-fold molar excess of acceptor dye and incubating both dyes with the TM3/4 sample for 3 h in total at room temperature before purification via size exclusion chromatography. Fluorescently labeled TM3/4 variants were reconstituted in freshly prepared POPC LUVs (16:0–18:1 PC, Avanti Polar Lipids, USA) with a diameter of ~110 nm as previously reported[3].

**Preparation and performance of automated multiwell plate measurements**

Prior to an automated multiwell-plate measurement, each well of a 96-well plate (Table S1) was passivated with 250 µL of a 1 mg/mL BSA solution (in phosphate buffered saline) for 5 min with subsequent removal of the BSA solution after incubation. This step was done to avoid nonspecific adsorption of the biomolecules on the chamber surface. In the case of S6, where high concentrations of GdmCl were used, we added 0.0005% (v/v) Tween 20 to the buffer instead of passivating with BSA (Tab. S5). For measurements of concentration gradients, as done for $hpT_5$ and S6, we prepared several stock solutions containing ~100 pM sample (labeled protein or oligo), buffer and different concentrations of salt/denaturant. A specific concentration per well was then obtained by mixing two stocks in a given ratio. For more complex experiments (*e.g.*, $dT_{70}$ in presence of SSB and RecA), multiple pipetting steps per well were necessary. For CFTR TM3/4 measurements, both corrector drugs, Lumacaftor (Lot number: 06855, purity: 99.06%, MedChemExpress LLC, USA) and Galicaftor (Lot number: S853501, purity: 98.93%, Selleck Chemicals LLC, USA), were dissolved in DMSO and stored at -20°C until use. For drug titration to the dialyzed TM3/4 variants in LUVs, a 2 mM stock solution of either Lumacaftor or Galicaftor in 50 mM Tris, pH 7.4 was prepared by adding the drug from DMSO into buffer followed by vortexing at full speed for 10 s and 5 min of sonication. Table S5 summarizes the experimental buffers and prepared stocks.

After passivation and the preparation of stocks, a pipetting time of usually 2-3 h was necessary to load a 96-well plate. To prevent evaporation of the solution during well plate preparation, we avoided volumes smaller than 20 µL and kept the well plate on ice during pipetting. In a test run with a liquid dispensing device (MANTIS®, FORMULATRIX, Bedford, MA, USA), which enables quick loading of 96-well plates by microdiaphragm pumps, the preparation time was reduced to less than 15 min. At the end of the preparation, we sealed the well plate with parafilm to prevent evaporation and concentration changes by loss of solvent over time. Further, condensation on the parafilm was avoided by heating it slightly above room temperature with a heating pad (Table S1). During the ~32 h measurement time (96 × 20 min), the immersion water of the objective was refreshed every 100 min for 5 s by a liquid dispenser system and the focus was kept 30 µm in solution using the Nikon Perfect Focus System (Table S1).

**Burst search algorithm and calculation of MFD parameter**

Single-molecule events were identified from the recorded photon stream using a burst search algorithm. This algorithm defines single-molecule events as such if more than $N_{\mathrm{ph}}$ adjacent photons exhibit an inter-photon time smaller than $ITP_{\mathrm{Burst}}$, after applying a Lee filter of four[4]. Fluorescence background was estimated from the photon stream by regions of 50 adjacent photons with a minimum inter-photon time of $ITP_{\mathrm{BG}}$. The three necessary parameters for the burst search algorithm are usually in the ranges of $30 < N_{\mathrm{ph}} < 200$, $5\ \mu\mathrm{s} < ITP_{\mathrm{Burst}} < 30\ \mu\mathrm{s}$, and $5\ \mu\mathrm{s} < ITP_{\mathrm{BG}} < 30\ \mu\mathrm{s}$. Note, that for measurements of high burst density, the background was estimated differently by selected regions of the time trace. Details are given in Tab. S6.

According to the color (*i.e.*, green (G) for donor and red (R) for acceptor fluorescence detection channel) and the microtime (emission delay time) of the photons, three different intensities were extracted for each fluorescence burst: the background subtracted fluorescence intensities in the donor and the acceptor channel after donor excitation $F_{\mathrm{GG}}$ and $F_{\mathrm{GR}}$, respectively, and in the acceptor channel after acceptor excitation $F_{\mathrm{RR}}$. The three intensities allowed then the calculation of the apparent/corrected stoichiometry and FRET efficiency values, $S_{\mathrm{app}}$, $E_{\mathrm{app}}$, $S$ and $E$, respectively, as follows[5]:

$$S_{\mathrm{app}} = \frac{F_{\mathrm{GR}} + F_{\mathrm{GG}}}{F_{\mathrm{GR}} + F_{\mathrm{GG}} + F_{\mathrm{RR}}} \tag{1}$$

$$E_{\mathrm{app}} = \frac{F_{\mathrm{GR}}}{F_{\mathrm{GR}} + F_{\mathrm{GG}}} \tag{2}$$

$$S = \frac{F_{\mathrm{GR}} - \alpha \cdot F_{\mathrm{RR}} - \beta \cdot F_{\mathrm{GG}} + \gamma \cdot F_{\mathrm{GG}}}{F_{\mathrm{GR}} - \alpha \cdot F_{\mathrm{RR}} - \beta \cdot F_{\mathrm{GG}} + \gamma \cdot F_{\mathrm{GG}} + F_{\mathrm{RR}}} \tag{3}$$

$$E = \frac{F_{\mathrm{GR}} - \alpha \cdot F_{\mathrm{RR}} - \beta \cdot F_{\mathrm{GG}}}{F_{\mathrm{GR}} - \alpha \cdot F_{\mathrm{RR}} - \beta \cdot F_{\mathrm{GG}} + \gamma \cdot F_{\mathrm{GG}}} \tag{4}$$

where $\alpha$, $\beta$, and $\gamma$, denote the correction factors for direct excitation, spectral crosstalk, and differences in detection efficiency and quantum yield (Tab. S6). In the cases, where two FRET populations were present at low and high FRET efficiency (open/unfolded and closed/folded conformation), the correction factors were directly extracted from the measurement (following Hellenkamp et al.[6]). In order to probe for fluorescence quenching and ultrafast conformational dynamics, the fluorescence lifetime of the donor and acceptor, $\tau_{\mathrm{D}}$ and $\tau_{\mathrm{A}}$, respectively, were calculated for each molecule from photon delay times (microtimes). Since the fluorescence background was in all measurements sufficiently low (< 5%)[7], the lifetimes were estimated by

$$\tau \approx \frac{1}{N}\sum_{i} t_{\mathrm{i}} - \langle t_{\mathrm{IRF}} \rangle \tag{5}$$

Here, the first term represents the average microtime of the respective color channel and time window (GG & GR or RR), and the second term the average microtime of the corresponding detector instrumental response function (IRF).

An important necessity for accurate FRET efficiencies is the free rotation of the fluorophore dipoles, which allows the assumption of $\kappa^2 = 2/3$ for the relative dipole orientations. The rotational freedom is encoded in the fluorescence anisotropy $r$. The anisotropy of donor and acceptor for each molecule was calculated using Eq. 6.

$$r = \frac{G \cdot F_{\parallel} - F_{\perp}}{G \cdot F_{\parallel} + 2F_{\perp}} \tag{6}$$

The polarization beam splitter of the setup was used to separate the parallel and perpendicular part of the fluorescence intensity, $F_{\parallel}$ and $F_{\perp}$, respectively, taking into account the difference in detection efficiency by the correction factor $G$.

The kernel-density based scores ALEX-2CDE and FRET-2CDE were derived for each molecule as previously described[8,9] using a characteristic kernel decay time of 75 µs and 12.5 µs, respectively.

**Data analysis of the 9- and 21-bp DNA ruler**

The FRET efficiency correction factors $\alpha$, $\beta$, and $\gamma$ were determined individually for each well (Fig. S4B). Therefore, we applied certain filter criteria for the estimation of $\alpha$: ALEX-2CDE > 10, FRET-2CDE < 10, and a minimum number of 10 photons after red excitation; for $\beta$: ALEX-2CDE > 10, FRET-2CDE < 10, $E_{app} > 0.8$, and a minimum number of 30 photons after green excitation; and for $\gamma$: ALEX-2CDE < 5, FRET-2CDE

< 10, $0.2 < S_{app} < 0.8$. The correction factor for direct excitation was derived by $\alpha = \langle S_{app}\rangle/(1 - \langle S_{app}\rangle)$, where $\langle S_{app}\rangle$ represents the average apparent stoichiometry of the filtered molecules. For the correction factor of the spectral crosstalk the average apparent FRET efficiency $\langle E_{app}\rangle$ of the filtered molecules was calculated instead and plugged into $\beta = \langle E_{app}\rangle/(1 - \langle E_{app}\rangle)$. Subsequently, the $\alpha$ and $\beta$ corrected stoichiometries ($S^*$) and FRET efficiencies ($E^*$) were calculated for the filtered molecules yielding two populations of 9 and 21bp molecules. The center positions of the two populations $\{1/\langle S^*\rangle_1, \langle E^*\rangle_1\}$ and $\{\{1/\langle S^*\rangle_2, \langle E^*\rangle_2\}\}$ were then fitted by a linear fit allowing the quantification of the detection correction factor by $\gamma = (n_{SE} - 1)/(n_{SE} - 1 + m_{SE})$. Here $n_{SE}$ and $m_{SE}$ denote the intercept and slope of the linear fit. The resulting 96 values (Fig. S4B) showed only minor variations of $\Delta\alpha/\alpha = 5.20\%$, $\Delta\beta/\beta = 5.62\%$, and $\Delta\gamma/\gamma = 6.47\%$. For comparison we calculated the theoretical correction factors using the manufacturing information of the emission/absorption spectra and quantum yields of the fluorophores (ATTO-TEC), as well as the transmission spectra of the dichroics and bandpass filters (Chroma and Semrock) and the photon detection efficiencies of the detectors (Excelitas). The theoretical values (black dashed lines, Fig. S4B) are in good agreement with the average measured correction factors.

For smFRET analysis, only double labeled molecules with both fluorophores active were extracted by demanding an ALEX-2CDE score smaller than 15 and a stoichiometry value between 0.25 and 0.75. Yielding an average count of $N = 1322 \pm 108$ molecules/well, only a minor decay of 206 number of molecules in the wells was observed for the 32 h of measurement (Fig. S4A).

The corrected FRET efficiencies (see Tab. S6) were sorted into histograms with bin size 0.03 and fitted by two Gaussian distributions (Fig. S4C). The resulting standard deviations, average FRET efficiencies, and 21 bp molecule fractions (Fig. S4D) of the 96 repeats allowed us to quantify the measurement precision and accuracy. To this end, we modeled the dye positions around the static DNA construct with accessible volume simulations[10] applying the recommended geometry parameters for ATTO 532 and ATTO 647N and the corresponding Förster radius of $R_0 = (5.06 \pm 0.05)$ nm. Latter was determined in a separate measurement and is in good agreement with the previously reported 5.1 nm[11]. To gain estimations for the prediction uncertainty we took the standard deviation of the Förster radius into account. The theoretical values are $E_{\text{9bp}}^{\text{theo}} = 0.777 \pm 0.009$ and $E_{\text{21bp}}^{\text{theo}} = 0.159 \pm 0.008$ as indicated in Fig. S4D (red and dashed lines). Using the predicted and 96 extracted average FRET efficiencies, $\langle E \rangle_{\text{AV}}$ and $\langle E \rangle_i$, respectively, we then assessed the measurement accuracy for the 9- and 21-bp construct by the mean absolute deviation $MAD = \sum_{i=1}^{N} |\langle E \rangle_{\text{AV}} - \langle E \rangle_i| \,/N$ resulting in $MAD(\text{9bp}) = 2.59\%$ and $MAD(\text{21bp}) = 8.20\%$. The precision (*i.e.*, repeatability), on the contrary, was assessed for the two constructs by the variance of the individual $\langle E \rangle_i$ values yielding $\sigma_{\langle E \rangle}(9bp) = 0.0052$ and $\sigma_{\langle E \rangle}(21bp) = 0.0049$. For the sake of completeness, the standard deviations of the Gaussian distributions are plotted in Fig. S4E.

**Analysis of the DNA hairpin hpT$_5$ opening and closing**

For the analysis of the DNA hairpin measurement, where we aimed to quantify the interconversion dynamics, we chose an alternative burst search algorithm. To this end, the photon stream was converted into an intensity time trace with bin time $T_{\text{b}} = 1\ \text{ms}$, and fluorescence bursts were identified having an intensity bigger than 100 kHz (photons/ms). Additionally, we demanded an ALEX-2CDE score smaller than 15 and an apparent stoichiometry between 0.3 and 0.7 to gain solely molecules with donor and acceptor active. For the remaining molecules, apparent FRET efficiency histograms were calculated from the background corrected donor and acceptor intensities $F_{\text{GG}}$ and $F_{\text{GR}}$ after donor excitation. Although the alternative burst search algorithm misses some molecules with burst durations smaller than $T_{\text{b}}$, the extracted FRET efficiency histograms exhibit reduced shot noise due to lack of convolution of the shot noise with the burst duration distribution. This leads to a higher contrast of dynamic features and a more accurate extraction of kinetic rates.

In a next step, we plotted the FRET-2CDE score over $E_{\text{app}}$ as shown in Fig. S5A. Here an arc of molecules becomes apparent with FRET-2CDE > 20 connecting the open and closed conformation of the DNA hairpin. These events originate from molecules undergoing conformational changes during the diffusion through the confocal volume, *i.e.*, millisecond dynamics. Millisecond opening and closing dynamics were found at all salt conditions. In order to extract kinetic rates from the characteristic shape of a FRET efficiency histogram (FEH), we employed the three-Gaussian (3G) approximation (red line, Fig. S5B) as described by Gopich and Szabo[12,13]. Briefly, the

3G approximation to extract kinetic rates of an interconverting two-state system is analytically expressed as the sum of three Gaussian distributions:

$$FEH(E) = A\sum_{i=0}^{2} c_{\mathrm{i}}(2\pi\sigma_i{}^2)^{-1/2} exp\left(-\frac{(E-\varepsilon_i)^2}{2\sigma_i{}^2}\right) \quad (7)$$

While the two Gaussians with subscripts $i = 1$ or $2$ refer to molecules in the folded and unfolded state, respectively, the Gaussian with subscript $i = 0$ represents molecules that appear at FRET efficiencies between the two states and undergo conformational changes (FRET-2CDE > 20). While $\varepsilon_1$ and $\varepsilon_2$ denote the FRET efficiencies of the folded and unfolded state, respectively, $A$ represents the total number of molecules sorted into the measured histogram. The remaining parameters ($c_{\mathrm{i}}$, $\sigma_{\mathrm{i}}{}^2$) describe the relative amplitudes and the variance of the distributions and are functions of $\varepsilon_1$ and $\varepsilon_2$ and the closing and opening rates $k_1$ and $k_2$. These parameters are given as following:

$$c_{\mathrm{i}} = p_{\mathrm{i}} exp(-k_{\mathrm{i}} T_{\mathrm{b}}) \qquad \mathrm{i} = 1{,}2$$

$$c_0 = 1 - c_1 - c_2$$

$$\sigma_i{}^2 = \varepsilon_{\mathrm{i}}(1-\varepsilon_{\mathrm{i}})\langle N^{-1}\rangle \qquad \mathrm{i} = 1{,}2$$

$$c_0\varepsilon_0 = \sum_{\mathrm{i}=1}^{2}(p_{\mathrm{i}} - c_{\mathrm{i}})\varepsilon_{\mathrm{i}} \quad (8)$$

$$c_0\sigma_0{}^2 = \langle\varepsilon\rangle_{\mathrm{eq}}\left(1-\langle\varepsilon\rangle_{\mathrm{eq}}\right)\langle N^{-1}\rangle +$$

$$+\,2p_1p_2(\varepsilon_2-\varepsilon_1)^2(kT_{\mathrm{b}} + e^{-kT_{\mathrm{b}}} - 1)(1-\langle N^{-1}\rangle)/(kT_{\mathrm{b}})^2 +$$

$$+\,\langle\varepsilon\rangle_{\mathrm{eq}}{}^2 - \sum_{\mathrm{i}=0}^{2} c_{\mathrm{i}}\varepsilon_{\mathrm{i}}{}^2 - \sum_{\mathrm{i}=1}^{2} c_{\mathrm{i}}\sigma_{\mathrm{i}}{}^2$$

Here, $p_{\mathrm{i}} = k_{3-\mathrm{i}}/k$ is the equilibrium probability of state i, $\langle\varepsilon\rangle_{\mathrm{eq}} = p_1\varepsilon_1 + p_2\varepsilon_2$ is the equilibrium average FRET efficiency, $\langle N^{-1}\rangle$ is the average inverse total number of photons per bin time $T_{\mathrm{b}}$, and $k = k_1 + k_2$ is the relaxation rate. The value $\langle N^{-1}\rangle$ is extracted from the photon statistics of the measured fluorescence bursts. The only fitting parameters are $\varepsilon_1$, $\varepsilon_2$, $k_1$ and $k_2$ and are derived by minimization of the reduced chi-square calculated from the measured and predicted FEHs.

The salt dependency of the hpT5 opening rate follows a linear trend (Fig. 2F) with $m_{\mathrm{open}} = -(1.31 \pm 0.04)\ \mathrm{ms^{-1}M^{-1}}$ being the slope and $k^0_{\mathrm{open}} = (1.89 \pm 0.03)\ \mathrm{ms^{-1}}$ being the extrapolated rate at zero salt. To demonstrate the high accuracy of fitting parameters derived from a 96-well plate measurement, we assessed the variability of $m_{\mathrm{open}}$ and $k^0_{\mathrm{open}}$. Therefore, for each number of data points (*i.e.*, opening rates) 500 bootstrap iterations were performed, in which $N_{\mathrm{DP}}$ of the 75 extracted rates were randomly selected and fitted by a linear equation. Fig. S5C shows the resulting standard deviations as a function of $N_{\mathrm{DP}}$.

The closing rate, on the contrary, shows a deviation from the linear trend at lower salt concentrations. To shed light on the salt dependency of $k_{close}$, we calculated the apparent concentration of the hpT5 proximal strand $A$ around its complementary strand $\bar{A}$. Therefore, the concentration was estimated from the spherical volume spanned by the average distance $R_{\mathrm{A-\bar{A}}}$ between the two ends of the dT21 loop. The distance $R_{\mathrm{A-\bar{A}}}$ was calculated from the mean-square end-to-end distance $\langle R_{\mathrm{A-\bar{A}}}{}^2 \rangle = 2l_{\mathrm{p}}l_{\mathrm{c}} - 2l_{\mathrm{p}}\left(1 - e^{-l_{\mathrm{c}}/l_{\mathrm{p}}}\right)$ of a worm-like chain with contour length $l_{\mathrm{c}}$ and persistence length $l_{\mathrm{p}}$. While for the contour length of the dT21 loop a value of $l_{\mathrm{c}} = 14.2\ \mathrm{nm}$ was estimated (0.68 nm/nt), for the persistence length the well characterized Eq. 9 of Chen et al.[14] was employed with respective parameters for NaCl and a dT-DNA strand.

$$l_{\mathrm{p}}(I) = l_{\mathrm{p}}(\infty) + \frac{l_{\mathrm{p}}(0) - l_{\mathrm{p}}(\infty)}{(bI)^{n/2} + 1} \qquad (9)$$

Here, $l_{\mathrm{p}}(0)$ and $l_{\mathrm{p}}(\infty)$ denote the persistence length at zero and infinite amount of salt. The other parameters $I$, $b$ and $n$ represent the ionic strength, the charge screening efficiency, and the scaling exponent. The strong correlation of the closing rate with concentration [A] as shown in Fig. S5D is well described by a linear fit with slope $k'_{close} = (0.305 \pm 0.007) \cdot 10^6\ \mathrm{s^{-1}M^{-1}}$ and offset $k_{\mathrm{offset}} = (-0.31 \pm 0.02) \cdot 10^3\ \mathrm{s^{-1}}$ (red line). Due to a lower and upper limit of the persistence length of $l_p(\infty) = 0.75\ \mathrm{nm}$ and $l_p(0) = 2.09\ \mathrm{nm}$[14], respectively, we can predict the lower and upper boundary of the hpT5 closing rate to $0.022\ \mathrm{ms^{-1}} < k_{\mathrm{close}} < 1.02\ \mathrm{ms^{-1}}$. It is important to note that the offset $k_{\mathrm{offset}}$ of the linear fitting function $k_{\mathrm{close}} = k'_{close}[\mathrm{A}] + k_{\mathrm{offset}}$ discriminates the closing rate by a constant negative value, which is unexpected for pure association of the two complementary strands. Nevertheless, as reported earlier[15] one reason for the bias of kinetic rates originates from steric hindrance. Since $hpT_5$ contains also a 25 bp long distal stem adjacent to the annealing site, it is very likely that the offset is caused by an altered sampling volume and hindered diffusion of the complementary strand.

**Analysis of S6 stability and expansion of the unfolded state**

Detected fluorescence bursts were filtered according to their ALEX-2CDE score and stoichiometry (ALEX-2CDE < 8 and 0.25 < $S$ < 0.75) yielding only S6 molecules with both FRET reporters (donor and acceptor) available. For each GdmCl condition, the corrected FRET efficiencies were then calculated (Tab. S6), sorted into histograms, and fitted by a static double Gaussian function (static 2G, Fig. S6C) with individual $E$-values and shared population widths $\sigma$. Fig. S6A shows the resulting average FRET efficiency of the folded and unfolded conformation $E_{\mathrm{F}}$ and $E_{\mathrm{U}}$ (black rectangles and circles), respectively, depending on the denaturant concentration. The global standard deviations were determined to $\sigma_{\mathrm{F}} = 0.06 \pm 0.002$ and $\sigma_{\mathrm{U}} = 0.085 \pm 0.002$. For latter analysis of the unfolded S6 chain expansion, the FRET efficiencies were corrected ($\tilde{E}(\tilde{n}, \tilde{\Phi}_D) \rightarrow E(n, \Phi_D)$) for the change in refractive index with increasing GdmCl concentration using

$$E = \frac{1}{1 + \left(\frac{n}{\tilde{n}}\right)^4 \frac{\tilde{\Phi}_{\mathrm{D}}}{\Phi_{\mathrm{D}}}\left(\frac{1}{\tilde{E}} - 1\right)} \quad (10)$$

with $n = 1.3325$ and $\tilde{n}$ being the refractive index in absence (water at 25°C) and presence of GdmCl, respectively (red rectangles and circles). Since no significant change in the donor lifetime was observed for all denaturant conditions, we assumed equal donor quantum yields $\tilde{\Phi}_{\mathrm{D}} = \Phi_{\mathrm{D}}$. During the ~32 h of the 96-well plate measurement a drop in molecule abundance was observed around $[\mathrm{GdmCl}] \approx 3\ \mathrm{M}$ (Fig. S6B). This drop coincides with the unfolding transition, and is, therefore most likely caused by unfolded molecules sticking to the chamber surface.

To determine the midpoint $[\mathrm{GdmCl}]_{1/2}$, slope $m$ and thermodynamic stability $\Delta G_{\mathrm{H2O}} = m \cdot [\mathrm{GdmCl}]_{1/2}$ of the S6 folding transition, the fractions of unfolded molecules were fitted by the linear extrapolation method (LEM, red line, Fig. 2I):

$$f_U = \frac{e^{m([\mathrm{GdmCl}]-[\mathrm{GdmCl}]_{1/2})/RT}}{1+e^{m([\mathrm{GdmCl}]-[\mathrm{GdmCl}]_{1/2})/RT}} \tag{11}$$

The fit resulted in $[\mathrm{GdmCl}]_{1/2} = (3.16 \pm 0.01)\ \mathrm{M}$ , $m = (8.5 \pm 0.2)\ \mathrm{kJ\ mol^{-1}M^{-1}}$ and $\Delta G_{\mathrm{H2O}} = (26.7 \pm 0.7)\ \mathrm{kJ\ mol^{-1}}$. Again, we assessed the variability of the two fitting parameters $[\mathrm{GdmCl}]_{1/2}$ and $m$ depending on the number of data points (*i.e.*, fractions of unfolded molecules) (Fig. S6D, red dots). The whiskers indicate the fitting errors.

Besides the global folding transition, we also investigated the coil-to-globule (CG) transition of S6 by examining the expansion of the unfolded peptide chain at denaturant concentrations bigger than 2 M. Therefore, the radii of gyration $R_{\mathrm{G}}$ were derived from the refractive index corrected FRET efficiencies using the Sanchez model[16]. Briefly, the theory of Sanchez as expressed below takes inter-residue as well as excluded volume interactions into account[17].

$$P(R_{\mathrm{G}}, \varepsilon, R_{\mathrm{G},\theta}) = Z^{-1} R_{\mathrm{G}}^{6}\, exp\left(-\frac{7}{2}\frac{R_{\mathrm{G}}^{2}}{R_{\mathrm{G},\theta}^{2}} + \frac{1}{2}N\varepsilon\Phi - N\frac{1-\Phi}{\Phi}\ln(1-\Phi)\right)$$

$$P(r|R_{\mathrm{G}}) = \frac{1}{\delta \cdot R_{\mathrm{G}}}\left[3\left(\frac{r}{\delta \cdot R_{\mathrm{G}}}\right)^{2} - \frac{9}{4}\left(\frac{r}{\delta \cdot R_{\mathrm{G}}}\right)^{3} + \frac{3}{16}\left(\frac{r}{\delta \cdot R_{\mathrm{G}}}\right)^{5}\right] \text{ with } 0 \le r \le 2\delta{\cdot}R_{\mathrm{G}} \tag{12}$$

$$E = \int_{0}^{l_{\mathrm{c}}} \frac{1}{1+\left(\frac{r}{R_0}\right)^{6}} \int_{R_{\mathrm{c}}}^{\frac{l_{\mathrm{c}}}{2}} P(r|R_{\mathrm{G}}) P(R_{\mathrm{G}}, \varepsilon, R_{\mathrm{G},\theta})\, \mathrm{d}R_{\mathrm{G}}\, \mathrm{d}r$$

Here $\delta$, $N$, $\varepsilon$, $l_{\mathrm{c}}$ and $Z$ denote the correction factor ($\delta = \sqrt{5}$), the number of amino acids between the donor and acceptor labeling position, the mean interaction energy between amino acids, the contour length, and the normalization factor[18], respectively. The parameter $\Phi = (R_{\mathrm{C}}/R_{\mathrm{G}})^{3}$ represents the volume fraction of the chain with respect to the radius of the most compact conformation $R_{\mathrm{C}}$. The value for $R_{\mathrm{C}}$ was derived from the crystal structure of S6 (PDB: 1RIS) and was found to be 1.35 nm. $R_0 = 5.06\ \mathrm{nm}$ denotes the Förster radius. For the Θ-state the theoretical radius $R_{\mathrm{G},\theta} = 2.21\ \mathrm{nm}$ of a Gaussian chain was used as initial guess. After fitting the measured FRET efficiency of each denaturant condition using Eq. 12, the extracted fitting parameter $\varepsilon$ was converted into the average radius of gyration by

$$R_{\mathrm{G}} = \left( \int_{R_{\mathrm{c}}}^{\frac{l_{\mathrm{c}}}{2}} {r_{\mathrm{G}}}^2 \cdot P\left(r_{\mathrm{G}}, \varepsilon, R_{\mathrm{G},\theta}\right) \mathrm{d}r_{\mathrm{G}} \right)^{1/2} \tag{13}$$

The resulting values are shown in Fig. S6E and are well described by the empirical equation $R_{\mathrm{G}} = R_{\mathrm{C}}\left(1 + \rho \frac{K \cdot a}{1 + K \cdot a}\right)$ (red line), where $\rho = 2.2 \pm 0.6$ is the relative change in radius of gyration at high GdmCl activity[19] $a$ and $K = 1.5 \pm 0.1$ is the effective binding constant of GdmCl to the polypeptide monomers[20]. In the next step, we aimed for the actual dimension of the Θ-state. Therefore, the scaling exponent $v$ was calculated according to Eq. 14.

$$R_{\mathrm{G}} = \sqrt{\frac{2 l_{\mathrm{p}} b}{(2v + 1)(2v + 2)} N^{v}} \tag{14}$$

Since the Θ-state is in a range of conformational transition, we applied the persistence lengths $l_{\mathrm{p,unf}} = 0.4\ \mathrm{nm}$ and $l_{\mathrm{p,fold}} = 0.53\ \mathrm{nm}$ corresponding to the unfolded and folded peptide, respectively,[16] (Fig. S6F). From the average GdmCl concentration where the two extrapolated curves (red and blue line) reach a scaling exponent of $v = 1/2$ the radius of gyration of the Θ-state was obtained as $R_{\mathrm{G},\Theta} = (2.38 \pm 0.17)\ \mathrm{nm}$. Finally, the expansion factor was calculated using $\alpha = R_{\mathrm{G}}/R_{\mathrm{G},\Theta}$ (Fig. S6G). Interestingly, the CG transition between expansion factor 1 and $1 + (19/22)(R_{\mathrm{C}}^3/R_{\Theta}^3)$ appears below the actual unfolding midpoint at $a_{\mathrm{GdmCl}} = 1.73$.

**Analysis of the competitive and cooperative interaction of RecA and SSB with ssDNA**

For the 96+12-well plate measurement, where $dT_{70}$ was subjected to different concentrations of SSB, we requested an ALEX-2CDE score smaller than 7 and a stoichiometry between 0.15 and 0.62 to ensure both fluorescent reporters were functioning. For each of the 20 min long measurements, the corrected FRET efficiency histogram was then calculated from the remaining molecules using the appropriate correction factors (Tab. S6). Fig. S7 summarizes all histograms in a 2D representation, clearly visible are the two concentration regimes of the 65 and 35 binding mode. In order to characterize the transitions of the two binding modes, we fitted each $E$-histogram with two Gaussian functions and calculated the fraction of $SSB_{65}$ molecules from the relative area of the peak at $E \approx 0.8$ (Fig. 3C). As a binding model, we assumed a reversible sequential three-state transition $\mathrm{T} + m \cdot \mathrm{S} \underset{K_\mathrm{S}}{\leftrightarrow} \mathrm{TS}_m + n \cdot \mathrm{S} \underset{K_\mathrm{S}^*}{\leftrightarrow} \mathrm{TS}_{m+n}$ comprising the three states: unbound $dT_{70}$ (T), $dT_{70}$-$SSB_{65}$ ($TS_m$), and $dT_{70}$-$SSB_{35}$ ($TS_{m+n}$). The fraction of $SSB_{65}$ molecules is then expressed as:

$$f_{\mathrm{SSB65}} = \frac{1}{1 + \left(\frac{c_{\mathrm{S},1/2}}{[\mathrm{SSB}]}\right)^m + \left(\frac{[\mathrm{SSB}]}{c^*_{\mathrm{S},1/2}}\right)^n} \quad (15)$$

Where $m$ and $n$ are the Hill coefficients, and $c_{\mathrm{S},1/2} = \sqrt[m]{K_\mathrm{S}}$ and $c^*_{\mathrm{S},1/2} = \sqrt[n]{K^*_\mathrm{S}}$ are the concentration of half occupancy of the 65 and 35 binding mode, respectively. Fitting the measured fractions by Eq. 15 revealed $c_{S,1/2} = (278 \pm 1)\ \mathrm{pM}$, $c^*_{S,1/2} = (480 \pm 17)\ \mathrm{nM}$, $m = (2.12 \pm 0.07)$ and $n = (1.19 \pm 0.04)$ (Fig. 3C, blue line).

For the 96-well plate measurement, with RecA and SSB present in solution, we demanded an ALEX-2CDE score smaller than 12 and a stoichiometry between 0.15 and 0.62 to filter for molecules with both donor and acceptor active. The remaining molecules were then sorted into a 2D histogram according to their corrected FRET efficiency (Tab. S6) and well index (Fig. S8A and S8B). To analyze the influence of RecA on the SSB binding behavior, we quantified the fraction of the most abundant population at $E \approx 0.8$ by calculating $f(E > 0.4)$ (Fig. 3E). As demonstrated before (Fig. 3B), at low concentrations of RecA the conformation at $E \approx 0.8$ can be assigned to the $SSB_{65}$ binding mode. However, we also observed a significant shift of that population to lower FRET efficiencies in the presence of high RecA concentrations indicating a mixed conformation of RecA monomers and SSB tetramer on $dT_{70}$. Hence, we assumed a reversible two-pathway six-state reaction scheme (Fig. S9) containing the known empty $dT_{70}$ (T), RecA filament ($TR_p$), $SSB_{65}$ ($TS_m$) and $SSB_{35}$ ($TS_{m+n}$) states, as well as the two additional mixed conformations of RecA-SSB ($TR_qS_m$) and RecA-2xSSB ($TR_rS_{m+n}$). In this reaction scheme the values $K_\mathrm{S} = {c_{\mathrm{S},1/2}}^m$, $K^*_\mathrm{S} = {c^*_{\mathrm{S},1/2}}^n$, $K_\mathrm{R} = {c_{\mathrm{Rp},1/2}}^p$, $\alpha \cdot K_\mathrm{R} = {c_{\mathrm{Rq},1/2}}^q$, and $\alpha \cdot \beta \cdot K_\mathrm{R} = {c_{\mathrm{Rr},1/2}}^r$ denote the dissociation constants with the corresponding Hill coefficients $m$, $n$, $p$, $q$, and $r$. Since it was unclear where the mixed states $TR_qS_m$ and $TR_rS_{m+n}$ appear on the FRET axis, and thus what states contribute to $f(E > 0.4)$, we performed 75 independent fittings with varying state and probability configurations using Eq. 16 and the previously determined values $c_{\mathrm{S},1/2}$, $c^*_{\mathrm{S},1/2}$, $m$ and $n$.

$$f(\vec{s},\vec{p}) = \frac{\left(p_1 + p_4(u(\vec{s}) - 1)\right)\left(\frac{c_{\mathrm{S},1/2}}{[\mathrm{SSB}]}\right)^m + \left(p_2 + p_5(v(\vec{s}) - 1)\right) + \left(p_3 + p_6(w(\vec{s}) - 1)\right)\left(\frac{[\mathrm{SSB}]}{c^*_{\mathrm{S},1/2}}\right)^n}{u(\vec{s})\left(\frac{c_{\mathrm{S},1/2}}{[\mathrm{SSB}]}\right)^n + v(\vec{s}) + w(\vec{s})\left(\frac{[\mathrm{SSB}]}{c^*_{\mathrm{S},1/2}}\right)^m}$$

$$u(\vec{s}) = s_1 + s_4\left(\frac{c_{\mathrm{Rp},\frac{1}{2}}}{[\mathrm{RecA}]}\right)^p \qquad (16)$$

$$v(\vec{s}) = s_2 + s_5\left(\frac{c_{\mathrm{Rq},1/2}}{[\mathrm{RecA}]}\right)^q$$

$$w(\vec{s}) = s_3 + s_6\left(\frac{c_{\mathrm{Rr},1/2}}{[\mathrm{RecA}]}\right)^r$$

In Eq. 16 the state and probability configuration is specified in the Boolean vectors $\vec{s}$ and $\vec{p}$, respectively, where the row of each vector refers to the state as indexed in Fig. S9. Only states $i$ which are available for the reaction are $s_{\mathrm{i}} = 1$ otherwise zero. The same applies for $\vec{p}$, where only states which contribute to $f(E > 0.4)$ are set to one. The goodness of fit of each fitting was evaluated by the reduced chi-square ${\chi_r}^2$ taking the number of fitting parameters into account. The result of the 75 fitting iterations is plotted in Fig. S10 showing the smallest reduced chi-square of $\chi_r^2 = 0.108$ for the full reaction scheme, where $f(E > 0.4)$ describes the fraction of molecules being in state $TS_m$ and $TR_qS_m$ (green star). The corresponding half concentrations of occupancy are $c_{Rp,1/2} = (425\ \pm 91)\ \mathrm{nM}$, $c_{Rq,1/2} = (237\ \pm 76)\ \mathrm{nM}$, and $c_{Rr,1/2} = (278\ \pm 94)\ \mathrm{nM}$ with Hill coefficients $p = (4.9\ \pm 1.5)$, $q = (1.9\ \pm 1.5)$, and r= $(3.1\ \pm 1.6)$. The optimal curve of the fitting procedure is shown in Fig. 3E (blue line). Additionally to the smFRET data, we also inspected the apparent fluorescence anisotropy of the acceptor and found confirmation for the $SSB_{65}$ formation at $c_{S,1/2} = (278\ \pm 1)\ \mathrm{pM}$ and the RecA filament formation along dT70 at $c_{Rp,1/2} = (425\ \pm 91)\ \mathrm{nM}$ (Fig. S8C and S8D).

**CFTR TM3/4 data analysis**

Single-molecule bursts were identified as such by having more than 100 adjacent photons ($N_{\mathrm{GG}} + N_{\mathrm{GR}} + N_{\mathrm{RR}}$) with an inter-photon time smaller than 30 µs after a Lee filter of 4. Fluorescence background was estimated from the photon stream by regions of 50 adjacent photons with a minimum inter-photon time of 30 µs for Lumacaftor and

40 µs for Galicaftor measurements, respectively. For further analysis, only double labeled molecules with both fluorophores active were selected by satisfying a stoichiometry value of 0.2 < $S$ < 0.7 and a brightness ratio of ALEX-2CDE < 7. In order to exclude multiple molecule events (per vesicle and in the confocal volume), a minimum number of 15 acceptor photons after donor excitation and a maximum number of 300 acceptor photons after acceptor excitation was demanded. Further, measurement impurities and acceptor quenched fractions were excluded by requesting an acceptor fluorescence lifetime bigger than 2 ns.

For each remaining molecule the background corrected fluorescence counts $F_{\mathrm{GG}}$, $F_{\mathrm{GR}}$ $F_{\mathrm{RR}}$ were derived and the apparent FRET $E_{\mathrm{app}} = F_{\mathrm{GR}}/(F_{\mathrm{GG}} + F_{\mathrm{GR}})$ was calculated. The resulting $E_{\mathrm{app}}$ values were collected in histograms with a bin size of 0.033 and fitted with 2 Gaussian distributions to derive the fraction of molecules in the closed conformation (Fig. S11A, S11B and S11C). Based on the deviation of the fluorescence lifetime of the acceptor ($\tau_{\mathrm{A}}$), we discarded measurements with high Galicaftor concentrations (>500 µM) due to the compromised fluorescence intensities (Fig. S11E). The characteristic $EC_{50}$ values of the TM3/4 variant V232D were extracted for Lumacaftor and Galicaftor by fitting the closed fractions $f_{\mathrm{C}}$ to the following modified Hill equation:

$$f_{\mathrm{C}} = \frac{(f_{\mathrm{C,max}} - f_{\mathrm{C,min}})}{1 + \left(\frac{EC_{50}}{[\mathrm{D}]}\right)^{n}} + f_{\mathrm{C,min}} \tag{17}$$

where $n$ represents the shape factor of the transition and $f_{\mathrm{C,min}}$ and $f_{\mathrm{C,max}}$ are the minimum and maximum fraction of closed molecules at zero and infinite drug concentration $[\mathrm{D}]$, respectively.

### Supporting Figures

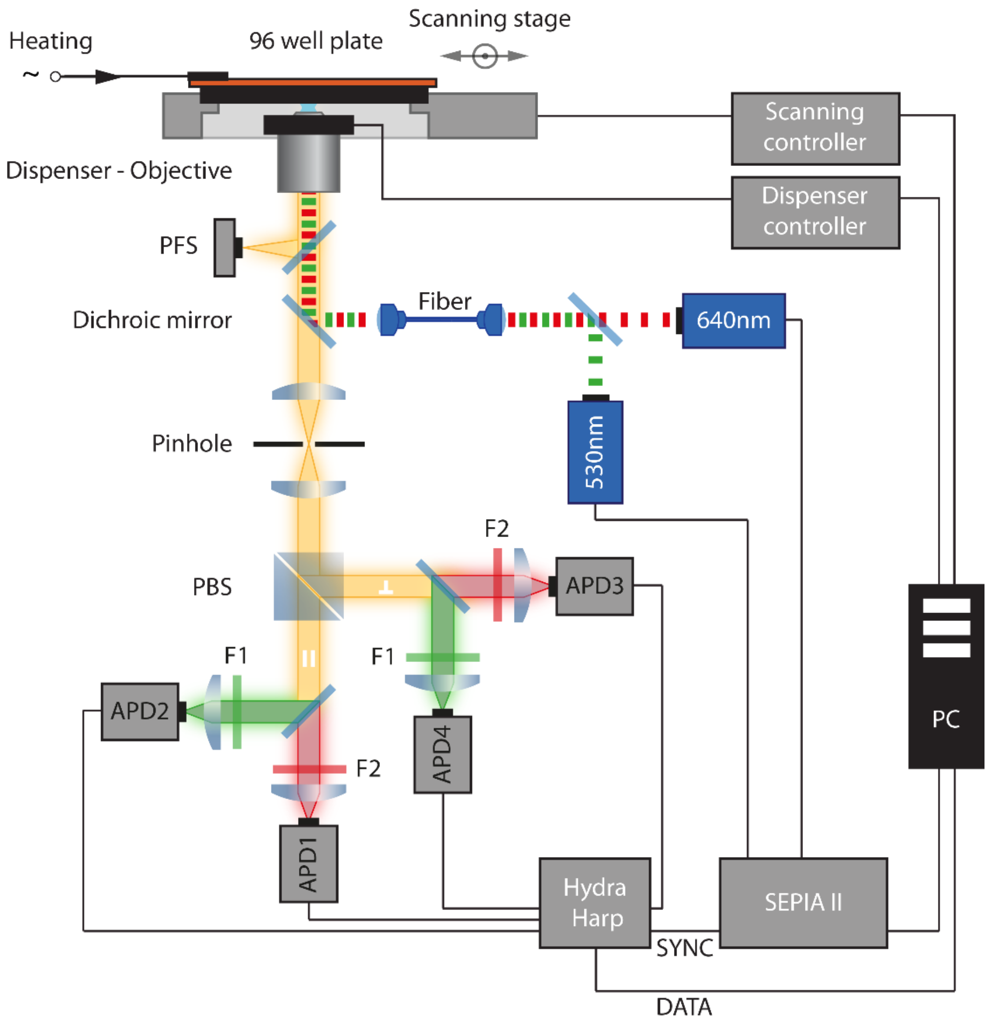


**Figure S1. Scheme of the custom-built automated multiwell plate, multi-parameter fluorescence detection setup for 96-well plate measurements.** The multiwell plate, multi-parameter fluorescence detection setup combines smFRET spectroscopy with time-correlated single photon counting (TCSPC) and fluorescence anisotropy detection. The motorized x-y stage mounted on the inverted microscope is capable of holding a 96-well plate and allows automated acquisition of 96 measurement conditions with almost zero dead time.

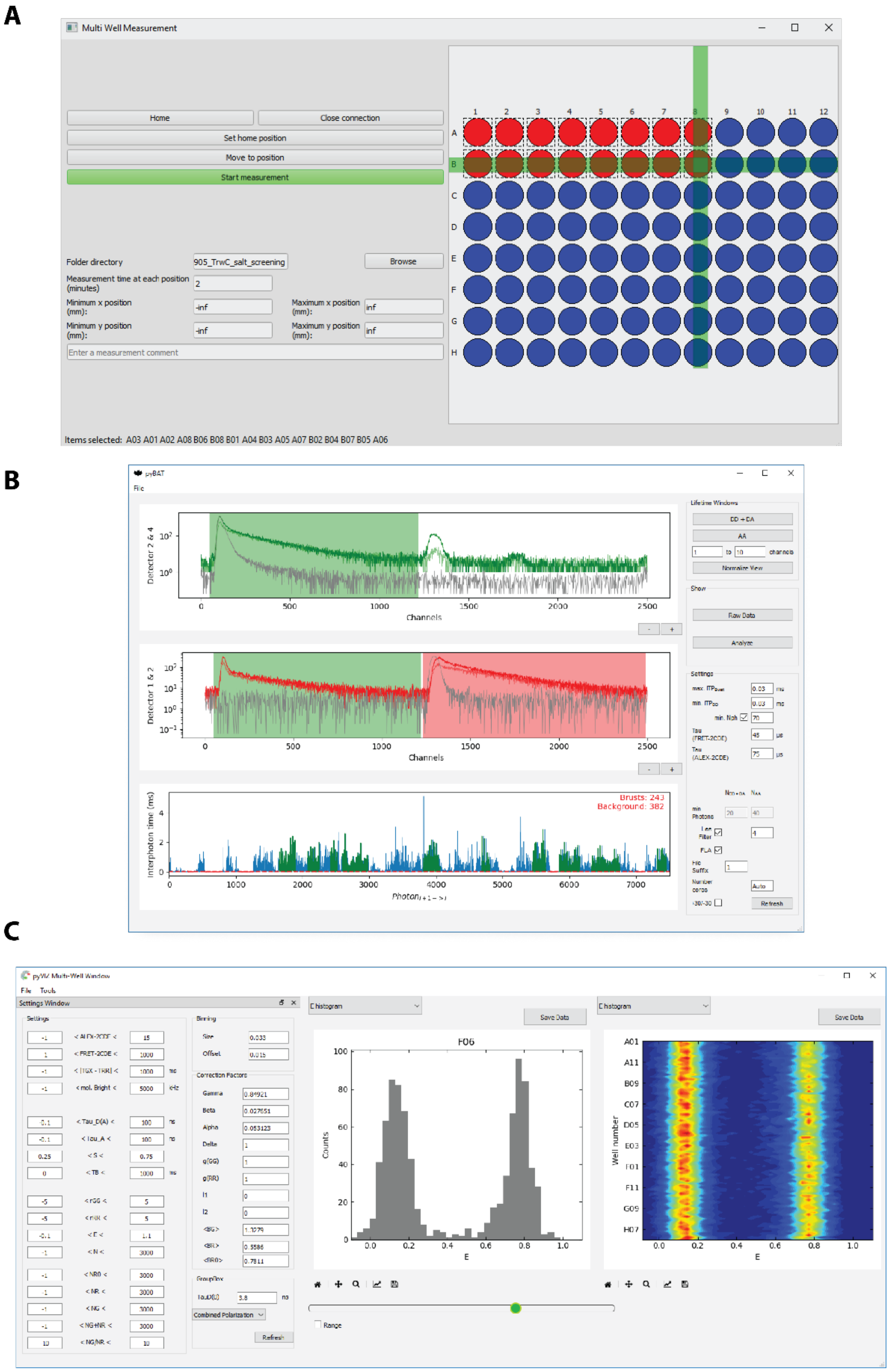


**Figure S2. Graphical user interfaces (GUI) of the autoFRET software package.** The software package consists of three parts: **(A)** pyMULTI, a GUI for hardware control of automated single and multiwell plate MFD measurements **(B)** pyBAT, a Burst Analysis Tool GUI for single and multiwell plate measurement data analysis, and **(C)** pyVIZ, a GUI for inspection of measurement results, data filtering, and data export.

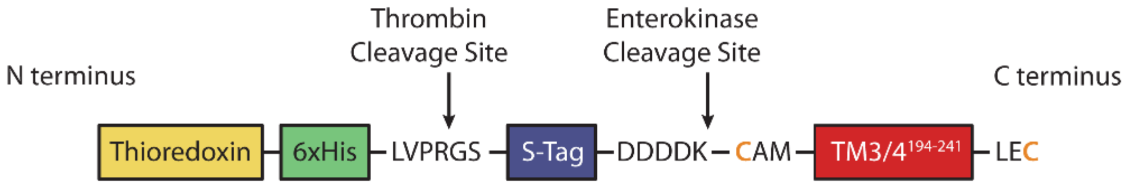


**Figure S3. Construct design of the TM3/4 fusion protein.** Thioredoxin (Trx, yellow) enables heat purification of the entire fusion construct, while a single His-tag (green) can be employed for purification via IMAC and reverse IMAC to remove non-digested protein and Trx after digestion by thrombin or enterokinase. The construct additionally bears an S-tag (violet) and two cysteine residues (orange) flanking the TM3/4 sequence (red, covering CFTR's residues 194 – 241).

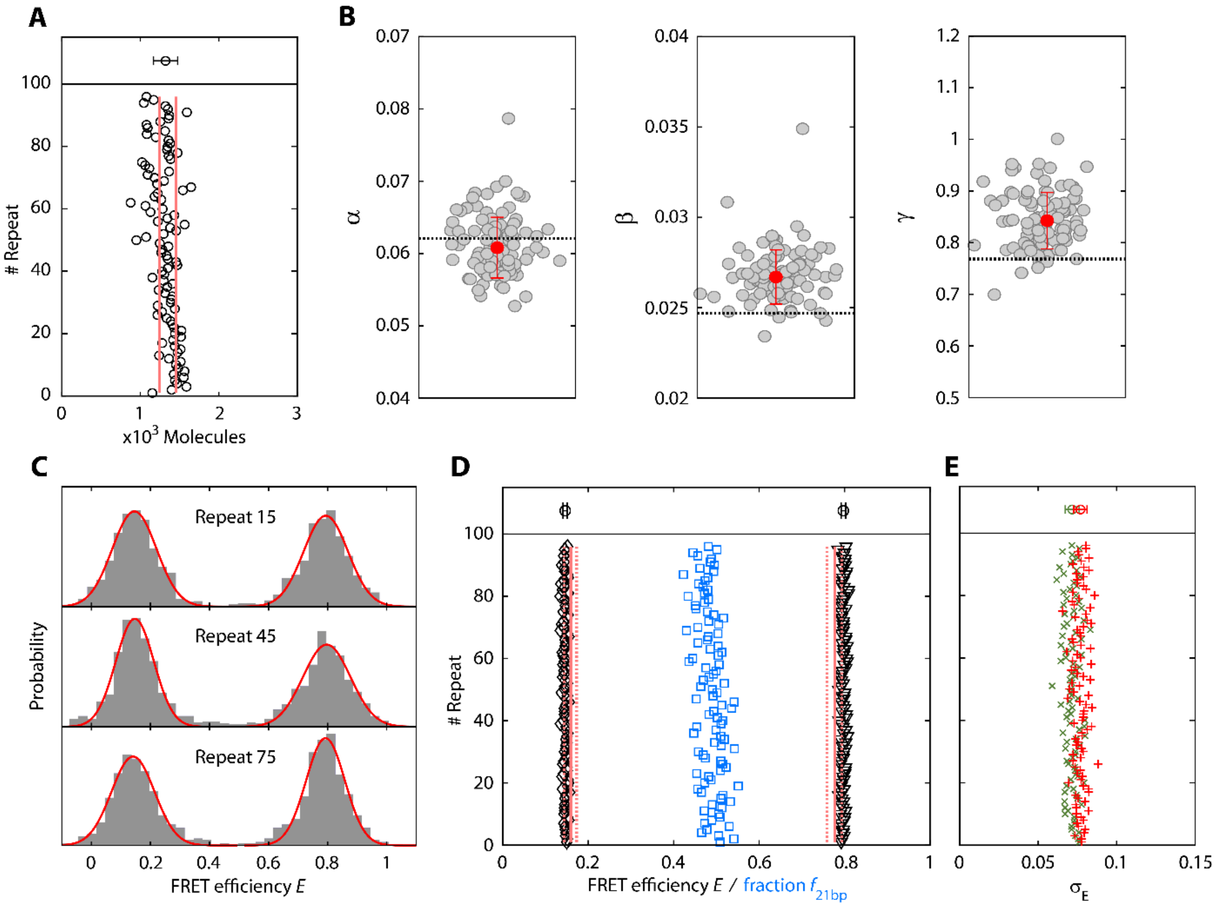


**Figure S4. Precision and accuracy of the automated multiwell plate approach with 9- and 21-bp DNA ruler substrates. (A)** Number of molecules per well remaining after filtering (circles lower panel) yielding an average count of $N = 1322 \pm 108$ molecules/well (upper panel) and only a minor decay of 206 molecules (red lines) over the 32 h of measurement time. **(B)** FRET efficiency correction factors were determined individually. The black dashed lines indicate the theoretical correction factors, which are in good agreement with the average measured correction factors (red dots; error bars are standard deviations). **(C)** For each well the FRET efficiencies of each burst were sorted into FRET efficiency histograms (grey bars) and fitted by two Gaussian distributions (red curves) to extract the center position, width and the molecule ratio of the FRET populations. **(D)** Average FRET efficiencies of the 9 and 21 bp FRET populations (triangles and diamonds, respectively) of all 96 repeats compared to the theoretical values of $E_{\text{9bp}}^{\text{theo}} = 0.777 \pm 0.009$ and $E_{\text{21bp}}^{\text{theo}} = 0.159 \pm 0.008$ (red lines) as derived from accessible volume simulations[10]. The dashed lines indicate the uncertainty of the prediction taking into account the Förster radius ($R_0 = (5.06 \pm 0.05)$ nm). The mean absolute deviation of the measured to the predicted values was smaller than 8.2% demonstrating the high accuracy of the multiwell plate measurements. We calculated mean FRET efficiencies of $\sigma_{\text{E9bp}} = 0.0052$ and $\sigma_{\text{E21bp}} = 0.0049$ together with a low standard deviation of the fractions of 21 bp molecules ($\langle f_{\text{21bp}} \rangle = 0.488$, blue rectangles) of $\sigma_{f\text{21bp}} = 0.027$ was observed. **(E)** Standard deviations of the 9 bp and 21 bp FRET populations (red pluses and green crosses, respectively) extracted from the widths of the fitted Gaussian distributions.

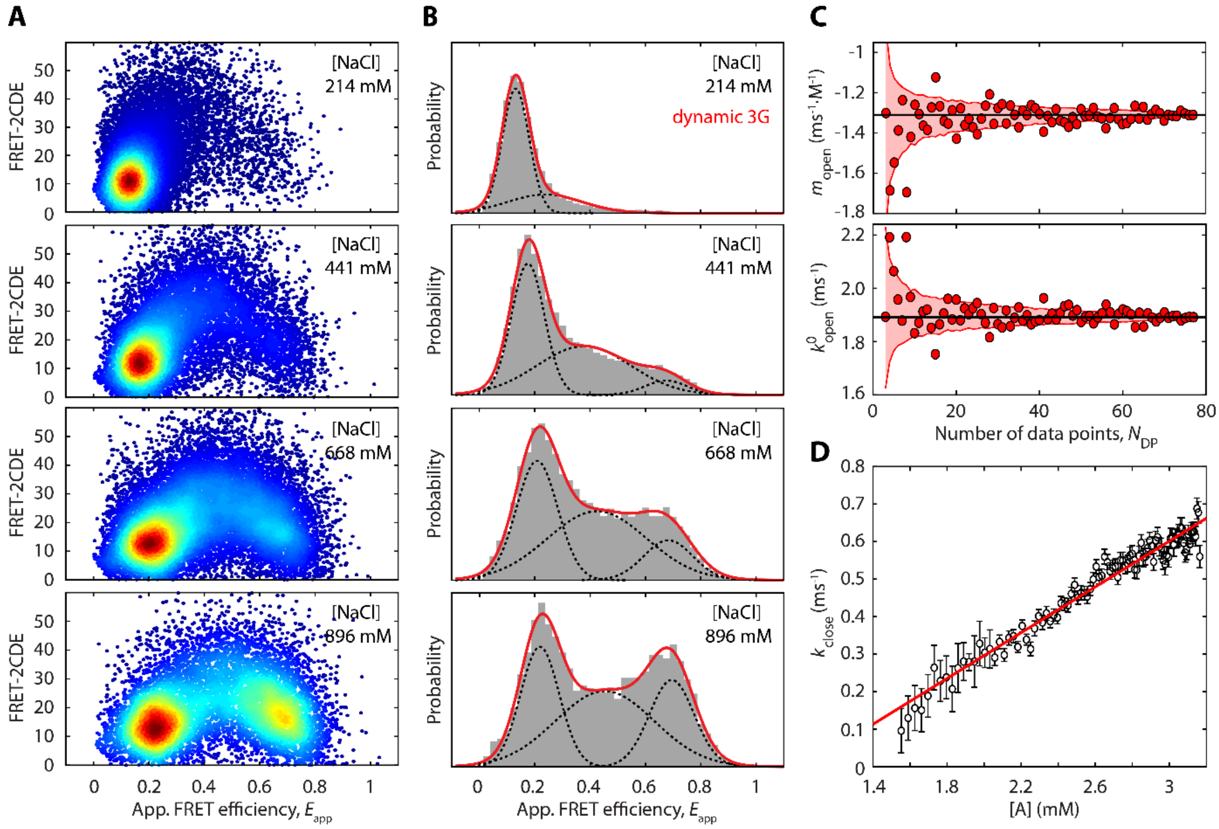


**Figure S5. Kinetic analysis of the millisecond opening and closing dynamics of hpT5. (A)** Identification of millisecond interconversion dynamics using the kernel density-based FRET-2CDE score[9]. The intermediate FRET population at $E_{\text{app}} \approx 0.5$ with FRET-2CDE > 20 originates from hpT5 molecules undergoing interconversion dynamics during diffusion through the confocal volume. **(B)** Apparent FRET efficiency histograms of hpT5 at different NaCl concentrations (grey bars). Kinetic rates of the millisecond opening and closing dynamics were extracted using the 3G approximation (dynamic 3G, solid red and dashed black lines)[8,12,13]. **(C)** Evaluation of the variability of fitting parameters (opening rates) calculated from increasing number of data points $N_{\text{DP}}$ (*i.e.*, kinetic rates). While the dots represent a single bootstrap sample, the area indicates the standard deviation derived from 500 bootstrap iterations. **(D)** Dependency of the measured closing rate $k_{close}$ on the calculated apparent concentration [A] of the hpT5 proximal strand $A$ around $\bar{A}$.

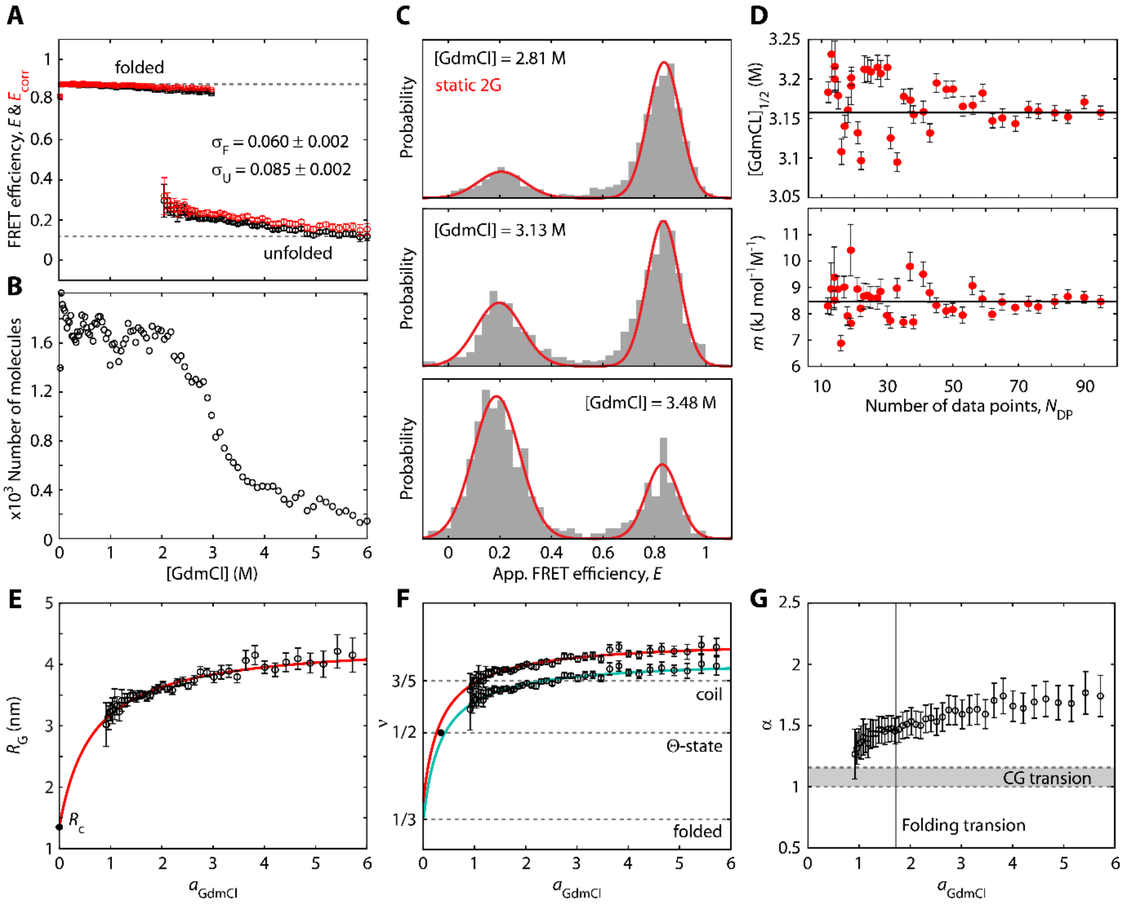


**Figure S6. Stability analysis and unfolded state expansion of S6. (A)** Average FRET efficiency of the folded and unfolded population of S6 at different GdmCl concentrations (black data points) extracted from 96 $E$-histograms. The FRET efficiencies were corrected for the change in refractive index with increasing denaturant concentration (red data points). **(B)** Number of extracted molecules of each 20-min long measurement at the respective denaturant concentration. **(C)** FRET efficiency histograms of S6 around the unfolding transition (grey bars). The fraction of unfolded molecules was quantified for each GdmCl concentration by a global double Gaussian fit with individual FRET efficiencies and shared widths. **(D)** Evaluation of the variability of the fitting parameters $[\mathrm{GdmCl}]_{1/2}$ and $m$ calculated from varying number of data points $N_{\mathrm{DP}}$ (*i.e.*, fraction of unfolded molecules). **(E-G)** Radius of gyration $R_{\mathrm{G}}$, scaling exponent $v$ and expansion factor $\alpha$ of the unfolded peptide chain of S6 as a function of GdmCl activity[19]. **(E)** The radius of gyration was calculated from the refractive index corrected FRET efficiencies of the unfolded state using the Sanchez model[21] including the most compact radius $R_{\mathrm{c}}$ at zero activity of GdmCl (black dot). The resulting values are well described by the empirical model of identical independent GdmCl binding sites[20] (red line, details in the SI). **(F)** The scaling exponent as converted from the radius of gyration of **(E)**. Two different persistence lengths were applied corresponding to the unfolded ($l_{\mathrm{p,unf}}$) and folded ($l_{\mathrm{p,fold}}$) peptide chain[16]. The radius of gyration of the Θ-state was obtained from the extrapolated curves as $R_{\mathrm{G},\Theta} = (2.38 \pm 0.17)\ \mathrm{nm}$. **(G)** The expansion factor calculated according to $\alpha = R_{\mathrm{G}}/R_{\mathrm{G},\Theta}$. The grey area indicates the coil-to-globule (CG) transition, while the grey line marks the unfolding midpoint.

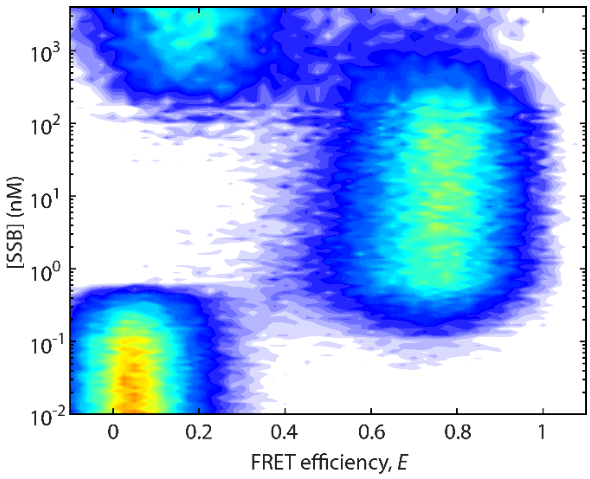


**Figure S7. 2D FRET efficiency histogram of dT$_{70}$ subjected to different SSB concentrations.**

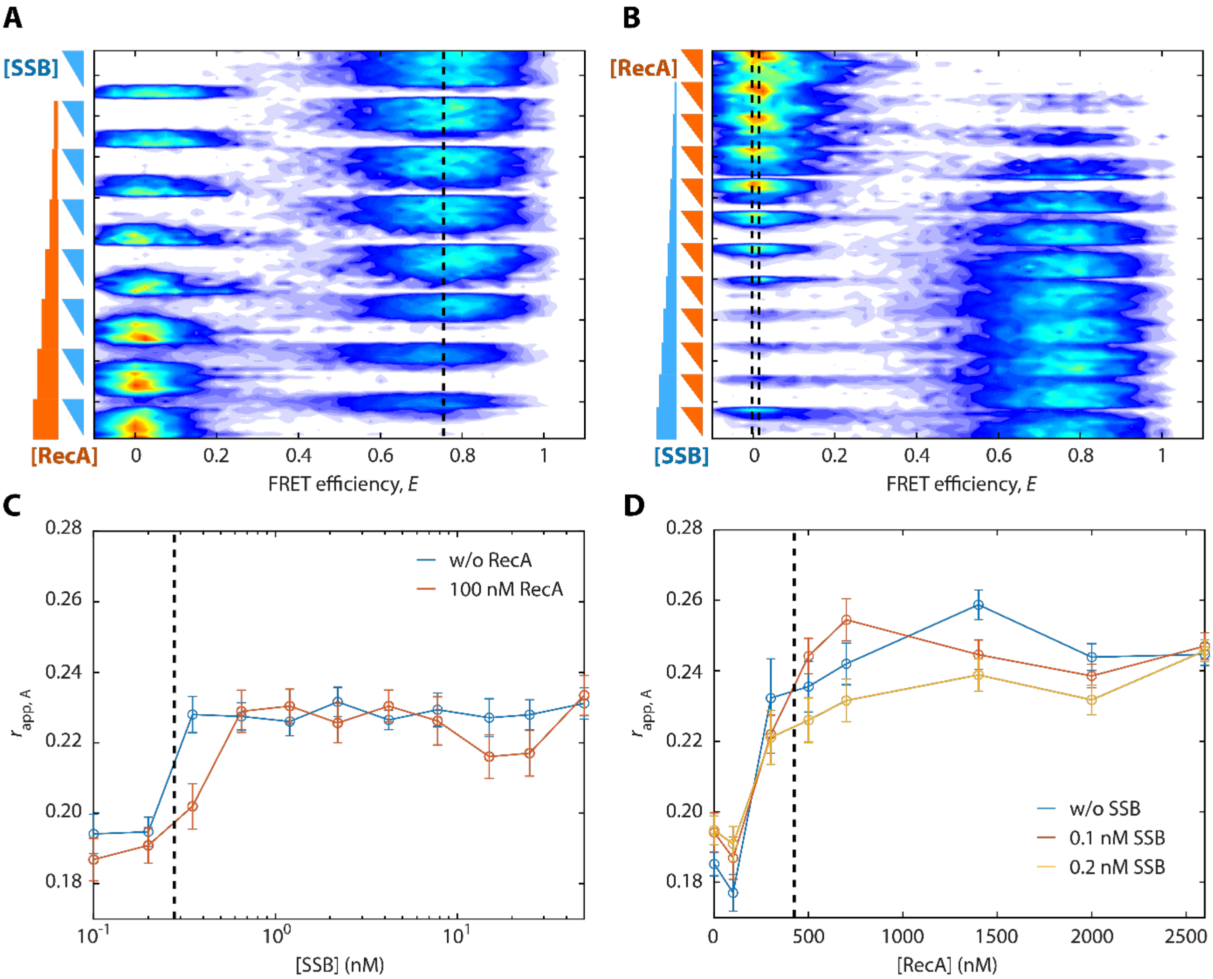


**Figure S8. 2D FRET efficiency histograms of the RecA-SSB-dT$_{70}$ experiment and fluorescence anisotropy of the acceptor. (A)** 2D FRET efficiency histogram from perspective of constant RecA but increasing SSB concentration. Dashed line indicates the center position of the SSB$_{65}$ population. **(B)** 2D FRET efficiency histogram of the same measurement from perspective of constant SSB but increasing RecA concentration. Dashed lines indicate the center positions of the dT$_{70}$ and RecA filament populations. **(C)** Apparent fluorescence anisotropy of the acceptor plotted against the SSB concentration in the absence and presence of 100 nM RecA. The transition at 300 pM agrees well with the formation of the SSB$_{65}$ binding mode at $c_{S,1/2} = (278 \pm 1)\,\mathrm{pM}$ (smFRET data). **(D)** Apparent fluorescence anisotropy of the acceptor plotted against RecA concentration in the absence and presence of low concentrations of SSB. The transition above 100 nM of RecA shows the start of filament formation on dT$_{70}$ and is consistent with the concentration of half occupancy of $c_{Rp,1/2} = (425 \pm 91)\,\mathrm{nM}$ derived from smFRET data.

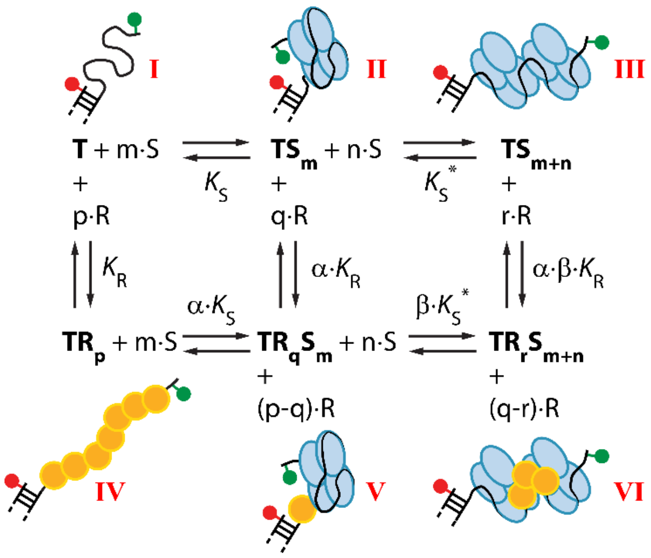


**Figure S9. Full reaction scheme describing the competitive and cooperative interaction of RecA and SSB with ssDNA.** 6-state model of the interactive binding of SSB and RecA to ssDNA, which contains the known $dT_{70}$ (T), RecA filament ($TR_p$), $SSB_{65}$ ($TS_m$) and $SSB_{35}$ ($TS_{m+n}$) states, as well as the two mixed states of RecA-SSB ($TR_qS_m$) and RecA-2xSSB ($TR_rS_{m+n}$). The states are numbered according to the Roman numerals. The values $K_S$, $K_S^*$, $K_R$, $\alpha \cdot K_R$, $\alpha \cdot \beta \cdot K_R$, $\alpha \cdot K_S$, $\beta \cdot K_S^*$ denote the respective dissociation constants with the corresponding Hill coefficients $m$, $n$, $p$, $q$, and $r$.

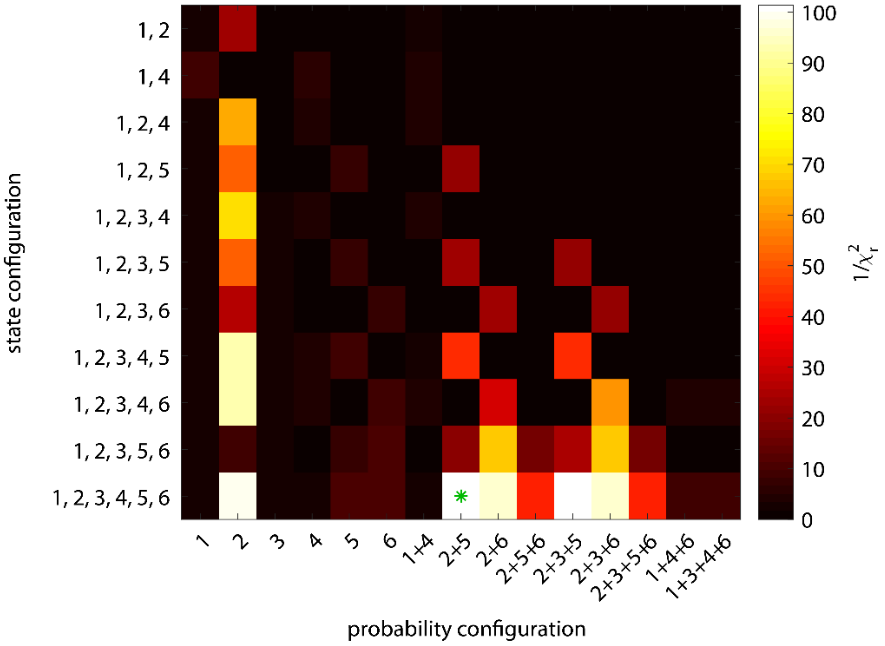


**Figure S10. Evaluation of different state and probability configurations.** The inverse reduced chi-squared is color-coded. The green star indicates the most likely state-probability-configuration with a reduced chi-squared of $\chi_r^2 = 0.108$.

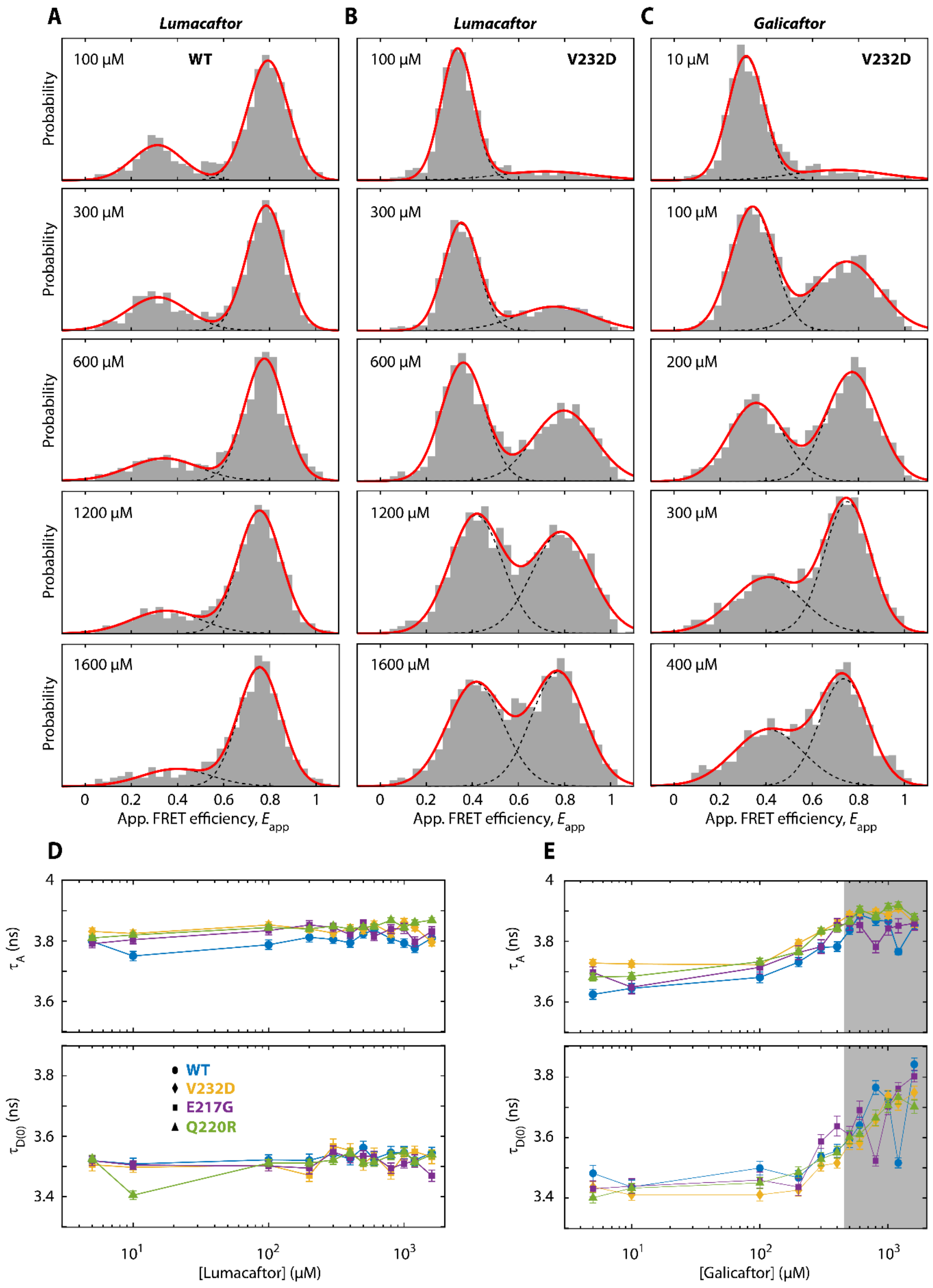


**Figure S11. Quantification of TM3/4 molecules in the closed conformation. (A)** Apparent FRET efficiency histograms of TM3/4 WT at different Lumacaftor (VX-809) concentrations (grey bars). The fraction of closed molecules was derived from a static two Gaussian fit (red and black dashed lines). **(B-C)** Lumacaftor and Galicaftor (ABBV-2222) response of TM3/4 mutant V232D. **(D-E)** Influence of the drug titration on acceptor and donor only fluorescence lifetimes $\tau_A$ and $\tau_{D(0)}$, respectively. While Lumacaftor shows no significant change in the average lifetime, Galicaftor alters the acceptor and donor lifetime at higher drug concentrations.

### Supporting Tables

**Table S1. Components used to establish the automated multiwell plate functionality in smFRET measurements**

| | | | |
|---|---|---|---|
| 1 | Motorized x-y stage for 96 well plate scanning | ASR100B120B-E03T3A with K0066 accessory kit | Zaber Technologies Inc., Vancouver, BC, Canada |
| 2 | Heating pad | ASIN B08JY4F81W | Lerway |
| 3 | Perfect focus unit | TI-ND6-PFS | Nikon, Tokyo, Japan |
| 4 | Liquid dispenser | Liquid dispenser, water immersion, 00-74-301-0000 | Märzhäuser Wetzlar GmbH & Co. KG, Wetzlar, Germany |
| 5 | 96-well plate | P96-1.5H-N, Glass Bottom Plates | IBL Baustoff+Labor GmbH, Gerasdorf, Vienna, Austria |

**Table S2. Runtime of pyBAT on different machines.**

| Processes (Threads) | Apple M1 Pro, 4 core CPU and 16 GB memory | Intel Xeon E3-1270, 4 core CPU and 32 GB memory | Intel i7-8700K, 6 core CPU and 32 GB memory |
|---|---|---|---|
| 1 | 24h | - | - |
| 7 | 3h | 13h | - |
| 11 | - | - | 7.3h |

**Table S3. Required python libraries.**

| Library | Source |
|---|---|
| Numpy | Harris, C.R., Millman, K.J., van der Walt, S.J. et al. *Array programming with NumPy*. Nature 585, 357–362 (2020). DOI: 10.1038/s41586-020-2649-2. |
| Pandas | McKinney, W. (2010, June). Data structures for statistical computing in python. In *Proceedings of the 9th Python in Science Conference* (Vol. 445, No. 1, pp. 51-56). |
| PyQT | PyQT (2012). PyQt Reference Guide. , . |
| Watchdog | https://github.com/gorakhargosh/watchdog |
| Numba | Lam, S. K., Pitrou, A., & Seibert, S. (2015). Numba: A llvm-based python jit compiler. In *Proceedings of the Second Workshop on the LLVM Compiler Infrastructure in HPC* (pp. 1–6). |
| Scipy | Virtanen, P., Gommers, R., Oliphant, T. E., Haberland, M., Reddy, T., Cournapeau, D., ... & Van Mulbregt, P. (2020). SciPy 1.0: fundamental algorithms for scientific computing in Python. *Nature methods*, *17*(3), 261-272. |
| Matplotlib | Hunter, J. D. (2007). Matplotlib: A 2D graphics environment. *Computing in science & engineering*, *9*(03), 90-95. |
| Joblib | https://github.com/joblib |
| Zaber Motion Library | https://gitlab.com/ZaberTech/zaber-motion-lib |
| pySerial | https://github.com/pyserial/pyserial |

**Table S4. Sequences of fluorescently labeled oligonucleotides and proteins used in this study.**

| | | |
|---|---|---|
| 1a | Top strand **DNA ruler** | Biotin - 5'- GCA TCA **X**CC AAG CGA CAC AAA CAG ACA ACC - 3' with X = dT - ATTO 532 |
| 1b | Bottom strand **DNA ruler 9bp** | 3' - CGT AGT AGG TTC GC**Y** GTG TTT GTC TGT TGG - 5' |
| 1c | Bottom strand **DNA ruler 21bp** | 3' - CGT AGT AGG TTC GCT GTG TTT GTC TG**Y** TGG - 5' with Y = dT - ATTO 647N |
| 2a | Top strand DNA hairpin **hpT$_5$** | 5' - ATTO 647N - TGG TT - (T)21 - AAC CAT TCT TCA CAA ACC AGT CCA AAC TAT CAA AAC TTA - 3' |
| 2b | Bottom strand DNA hairpin **hpT$_5$** | 3' - GTG T**Z**T GGT CAG GTT TGA TAG TTT TGA AT - 5' with Z = dT - ATTO 532 |
| 3a | FRET anchor **dT$_{70}$** | 5' - ATTO 647N - GCC TCG CTG CCG TCG CCA - biotin |
| 3b | ssDNA part **dT$_{70}$** | 5' - TGG CGA CGG CAG CGA GGC TTT TTT TTT TTT TTT TTT TTT TTT TTT TTT TTT TTT TTT TTT TTT TTT TTT TTT TTT TTT TTT TTT - ATTO 532 |
| 4 | **S6 M1C/F97C** | **C**RRYEVNIVL NPNLDQSQLA LEKEIIQRAL ENYGARVEKV EELGLRRLAY PIAKDPQGYF LWYQVEMPED RVNDLARELR IRDNVRRVMV VKSQEP**C**LAN A |
| 5a | **TM3/4 WT** | **C**AM - $^{194}$GLALAHFVWI APLQVALLMG LIWELLQASA FAGLGFLIVL ALFQAGLG$^{241}$ - LE**C** |
| 5b | **TM3/4 V232D** | **C**AM - $^{194}$GLALAHFVWI APLQVALLMG LIWELLQASA FAGLGFLI**D**L ALFQAGLG$^{241}$ - LE**C** |
| 5c | **TM3/4 E217G** | **C**AM - $^{194}$GLALAHFVWI APLQVALLMG LIW**G**LLQASA FAGLGFLIVL ALFQAGLG$^{241}$ – LE**C** |
| 5d | **TM3/4 Q220R** | **C**AM - $^{194}$GLALAHFVWI APLQVALLMG LIWELL**R**ASA FAGLGFLIVL ALFQAGLG$^{241}$ - LE**C** |

**Table S5. Measurement conditions and preparations.**

| Measurement | Buffer | Passivation | Preparation |
|---|---|---|---|
| DNA ruler | 20 mM Tris-HCl (pH 8), 50 mM NaCl | BSA | Pipetted from one stock solution containing ~100 pM DNA ruler in a 1:1 ratio of the 9 and 21 bp construct and buffer |
| DNA hairpin $hpT_5$ | 20 mM Tris-HCl (pH 8), 100 to 1000 mM NaCl | BSA | Pipetted from three stock solutions containing ~100pM $hpT_5$, buffer and different salt concentrations of 100, 500 and 1000 mM NaCl.<br><br>wells A01 to D12: different ratios of stock 1 (100 mM) and stock 2 (500 mM)<br><br>wells E01 to H12: different ratios of stock 2 (500 mM) and stock 3 (1000 mM) |
| S6 | 50 mM Tris-HCl pH 8, 150 mM NaCl, 2 mM TCEP, 0 to 6 M GdmCl | Tween20 in solution | Pipetted from 4 stock solutions containing ~100 pM S6, buffer and different denaturant concentration of 0, 3, 1.5 and 7.3 M GdmCl.<br><br>wells A01 to D12: different ratios of stock 1 (0 M) and stock 2 (3 M)<br><br>wells E01 to H12: different ratios of stock 3 (1.5 M) and stock 4 (7.3 M) |
| $dT_{70}$ + SSB | 50 mM Tris-acetate (pH 7.7), 5mM Mg-acetate, 50 mM Na-acetate | BSA | 96+12-well plate were loaded in 3 pipetting steps per well up to a final volume of 100 µL (~100 pM $dT_{70}$) using the following stocks:<br>- 0.5 nM $dT_{70}$<br>- buffer only<br>- SSB with 0.2, 2, 200, 2000 or 20000 nM |
| $dT_{70}$ + RecA + SSB | 50 mM Tris-acetate (pH 7.7), 5 mM Mg-acetate, 50 mM Na-acetate | BSA | 96-well plate was loaded in 4 pipetting steps per well up to a final volume of 100 µL (~100 pM $dT_{70}$) using the following stocks:<br>- 0.5 nM $dT_{70}$ and 80 mM ATP in buffer<br>- buffer only<br>- 2 stocks of RecA with 5 µM or and 52 µM<br>- 3 stocks of SSB with 2, 20, or 200 nM |
| TM3/4 variants | 50 mM Tris-HCl (pH 7.4) | BSA | To each well, 16 µL of dialyzed TM3/4 in LUVs was added and supplemented with the respective amount of drug stock to concentrations between 5 and 1600 µM to a total volume of 100 µL. |

**Table S6. Burst search parameters and FRET correction factors.**

| | *Parameters* | | | *Correction factors* | | |
|---|---|---|---|---|---|---|
| **Measurement** | $IPT_{\mathrm{Burst}}$ **(µs)** | $IPT_{\mathrm{BG}}$ **(µs)** | $N_{\mathrm{ph}}$ | $\alpha$ | $\beta$ | $\gamma$ |
| DNA ruler | 30 | 30 | 50 | Individual correction factors $\langle\alpha\rangle$=0.0608, $\langle\beta\rangle$=0.0267, $\langle\gamma\rangle$=0.8426 | | |
| DNA hairpin $hpT_5$ | 15 | 15 | 100 | 0 | 0 | 1 |
| S6 | 50 | 50 | 50 | 0.0854 | 0.0302 | 0.6601 |
| $dT_{70}$ + RecA + SSB | 50 | 50 | 40 | 0.0450 | 0.0805 | 0.9561 |
| TM3/4 variants | 30 | 30 (Lumacaftor)<br>40 (Galicaftor) | 100 | 0 | 0 | 1 |

**Table S7. NaCl concentration per well used for the DNA hairpin dynamic experiment**

| [NaCl] (mM) | 1 | 2 | 3 | 4 | 5 | 6 | 7 | 8 | 9 | 10 | 11 | 12 |
|---|---|---|---|---|---|---|---|---|---|---|---|---|
| **A** | 100.00 | 109.47 | 118.95 | 128.42 | 137.89 | 147.37 | 156.84 | 166.32 | 175.79 | 185.26 | 194.74 | 204.21 |
| **B** | 213.68 | 223.16 | 232.63 | 242.11 | 251.58 | 261.05 | 270.53 | 280.00 | 289.47 | 298.95 | 308.42 | 317.89 |
| **C** | 327.37 | 336.84 | 346.32 | 355.79 | 365.26 | 374.74 | 384.21 | 393.68 | 403.16 | 412.63 | 422.11 | 431.58 |
| **D** | 441.05 | 450.53 | 460.00 | 469.47 | 478.95 | 488.42 | 497.89 | 507.37 | 516.84 | 526.32 | 535.79 | 545.26 |
| **E** | 554.74 | 564.21 | 573.68 | 583.16 | 592.63 | 602.11 | 611.58 | 621.05 | 630.53 | 640.00 | 649.47 | 658.95 |
| **F** | 668.42 | 677.89 | 687.37 | 696.84 | 706.32 | 715.79 | 725.26 | 734.74 | 744.21 | 753.68 | 763.16 | 772.63 |
| **G** | 782.11 | 791.58 | 801.05 | 810.53 | 820.00 | 829.47 | 838.95 | 848.42 | 857.89 | 867.37 | 876.84 | 886.32 |
| **H** | 895.79 | 905.26 | 914.74 | 924.21 | 933.68 | 943.16 | 952.63 | 962.11 | 971.58 | 981.05 | 990.53 | 1000.00 |

**Table S8. GdmCl concentration per well used for the S6 unfolding experiment**

| [GdmCl] (M) | 1 | 2 | 3 | 4 | 5 | 6 | 7 | 8 | 9 | 10 | 11 | 12 |
|---|---|---|---|---|---|---|---|---|---|---|---|---|
| **A** | 0.020 | 0.041 | 0.063 | 0.085 | 0.107 | 0.129 | 0.152 | 0.176 | 0.200 | 0.225 | 0.250 | 0.275 |
| **B** | 0.302 | 0.328 | 0.355 | 0.383 | 0.411 | 0.440 | 0.470 | 0.500 | 0.531 | 0.562 | 0.594 | 0.627 |
| **C** | 0.660 | 0.694 | 0.729 | 0.764 | 0.800 | 0.837 | 0.875 | 0.913 | 0.952 | 0.992 | 1.033 | 1.074 |
| **D** | 1.117 | 1.160 | 1.205 | 1.250 | 1.296 | 1.343 | 1.391 | 1.440 | 1.490 | 1.541 | 1.593 | 1.646 |
| **E** | 1.700 | 1.755 | 1.812 | 1.869 | 1.928 | 1.988 | 2.049 | 2.112 | 2.175 | 2.240 | 2.307 | 2.374 |
| **F** | 2.443 | 2.514 | 2.586 | 2.659 | 2.734 | 2.811 | 2.889 | 2.968 | 3.050 | 3.133 | 3.217 | 3.303 |
| **G** | 3.392 | 3.482 | 3.573 | 3.667 | 3.763 | 3.860 | 3.959 | 4.061 | 4.165 | 4.271 | 4.378 | 4.489 |
| **H** | 4.601 | 4.716 | 4.833 | 4.952 | 5.074 | 5.198 | 5.325 | 5.455 | 5.587 | 5.722 | 5.860 | 6.000 |

**Table S9: SSB concentration in each well used to probe the two binding modes of SSB**

| [SSB] (nM) | 1 | 2 | 3 | 4 | 5 | 6 | 7 | 8 | 9 | 10 | 11 | 12 |
|---|---|---|---|---|---|---|---|---|---|---|---|---|
| **A** | 0.01 | 0.011 | 0.012 | 0.014 | 0.015 | 0.017 | 0.019 | 0.021 | 0.023 | 0.026 | 0.028 | 0.031 |
| **B** | 0.035 | 0.039 | 0.043 | 0.048 | 0.053 | 0.059 | 0.065 | 0.072 | 0.08 | 0.089 | 0.099 | 0.11 |
| **C** | 0.122 | 0.136 | 0.15 | 0.168 | 0.186 | 0.206 | 0.228 | 0.254 | 0.282 | 0.312 | 0.346 | 0.384 |
| **D** | 0.426 | 0.474 | 0.526 | 0.584 | 0.648 | 0.718 | 0.798 | 0.886 | 0.982 | 1.09 | 1.21 | 1.342 |
| **E** | 1.4 | 1.6 | 1.8 | 2 | 2.2 | 2.6 | 2.8 | 3 | 3.4 | 3.8 | 4.2 | 4.6 |
| **F** | 5.2 | 5.8 | 6.4 | 7.2 | 7.8 | 8.8 | 9.8 | 10.8 | 12 | 13.4 | 14.8 | 16.4 |
| **G** | 18.2 | 20.2 | 22.4 | 24.8 | 27.6 | 30.6 | 34 | 37.8 | 41.8 | 46.4 | 51.6 | 57.2 |
| **H** | 64 | 70 | 78 | 86 | 96 | 108 | 118 | 132 | 146 | 162 | 180 | 200 |

| [SSB] (nM) | 1 | 2 | 3 | 4 | 5 | 6 | 7 | 8 | 9 | 10 | 11 | 12 |
|---|---|---|---|---|---|---|---|---|---|---|---|---|
| **A** | 320 | 400 | 500 | 640 | 800 | 1000 | 1260 | 1600 | 2000 | 2520 | 3180 | 4000 |
| **B** | - | - | - | - | - | - | - | - | - | - | - | - |
| **C** | - | - | - | - | - | - | - | - | - | - | - | - |
| **D** | - | - | - | - | - | - | - | - | - | - | - | - |
| **E** | - | - | - | - | - | - | - | - | - | - | - | - |
| **F** | - | - | - | - | - | - | - | - | - | - | - | - |
| **G** | - | - | - | - | - | - | - | - | - | - | - | - |
| **H** | - | - | - | - | - | - | - | - | - | - | - | - |

**Table S10. RecA and SSB concentrations in each well used for the competitive/cooperative ssDNA binding experiment**

| | 1 | 2 | 3 | 4 | 5 | 6 | 7 | 8 | 9 | 10 | 11 | 12 |
|---|---|---|---|---|---|---|---|---|---|---|---|---|
| **A** | [RecA] =0μM [SSB] =0 | [RecA] =0μM [SSB] =0.1nM | [RecA] =0μM [SSB] =0.2nM | [RecA] =0μM [SSB] =0.35nM | [RecA] =0μM [SSB] =0.65nM | [RecA] =0μM [SSB] =1.2nM | [RecA] =0μM [SSB] =2.2nM | [RecA] =0μM [SSB] =4.2nM | [RecA] =0μM [SSB] =7.8nM | [RecA] =0μM [SSB] =15nM | [RecA] =0μM [SSB] =25nM | [RecA] =0μM [SSB] =50nM |
| **B** | [RecA] =0.1μM [SSB] =0 | [RecA] =0.1μM [SSB] =0.1nM | [RecA] =0.1μM [SSB] =0.2nM | [RecA] =0.1μM [SSB] =0.35nM | [RecA] =0.1μM [SSB] =0.65nM | [RecA] =0.1μM [SSB] =1.2nM | [RecA] =0.1μM [SSB] =2.2nM | [RecA] =0.1μM [SSB] =4.2nM | [RecA] =0.1μM [SSB] =7.8nM | [RecA] =0.1μM [SSB] =15nM | [RecA] =0.1μM [SSB] =25nM | [RecA] =0.1μM [SSB] =50nM |
| **C** | [RecA] =0.3μM [SSB] =0 | [RecA] =0.3μM [SSB] =0.1nM | [RecA] =0.3μM [SSB] =0.2nM | [RecA] =0.3μM [SSB] =0.35nM | [RecA] =0.3μM [SSB] =0.65nM | [RecA] =0.3μM [SSB] =1.2nM | [RecA] =0.3μM [SSB] =2.2nM | [RecA] =0.3μM [SSB] =4.2nM | [RecA] =0.3μM [SSB] =7.8nM | [RecA] =0.3μM [SSB] =15nM | [RecA] =0.3μM [SSB] =25nM | [RecA] =0.3μM [SSB] =50nM |
| **D** | [RecA] =0.5μM [SSB] =0 | [RecA] =0.5μM [SSB] =0.1nM | [RecA] =0.5μM [SSB] =0.2nM | [RecA] =0.5μM [SSB] =0.35nM | [RecA] =0.5μM [SSB] =0.65nM | [RecA] =0.5μM [SSB] =1.2nM | [RecA] =0.5μM [SSB] =2.2nM | [RecA] =0.5μM [SSB] =4.2nM | [RecA] =0.5μM [SSB] =7.8nM | [RecA] =0.5μM [SSB] =15nM | [RecA] =0.5μM [SSB] =25nM | [RecA] =0.5μM [SSB] =50nM |
| **E** | [RecA] =0.7μM [SSB] =0 | [RecA] =0.7μM [SSB] =0.1nM | [RecA] =0.7μM [SSB] =0.2nM | [RecA] =0.7μM [SSB] =0.35nM | [RecA] =0.7μM [SSB] =0.65nM | [RecA] =0.7μM [SSB] =1.2nM | [RecA] =0.7μM [SSB] =2.2nM | [RecA] =0.7μM [SSB] =4.2nM | [RecA] =0.7μM [SSB] =7.8nM | [RecA] =0.7μM [SSB] =15nM | [RecA] =0.7μM [SSB] =25nM | [RecA] =0.7μM [SSB] =50nM |
| **F** | [RecA] =1.4μM [SSB] =0 | [RecA] =1.4μM [SSB] =0.1nM | [RecA] =1.4μM [SSB] =0.2nM | [RecA] =1.4μM [SSB] =0.35nM | [RecA] =1.4μM [SSB] =0.65nM | [RecA] =1.4μM [SSB] =1.2nM | [RecA] =1.4μM [SSB] =2.2nM | [RecA] =1.4μM [SSB] =4.2nM | [RecA] =1.4μM [SSB] =7.8nM | [RecA] =1.4μM [SSB] =15nM | [RecA] =1.4μM [SSB] =25nM | [RecA] =1.4μM [SSB] =50nM |
| **G** | [RecA] =2.0μM [SSB] =0 | [RecA] =2.0μM [SSB] =0.1nM | [RecA] =2.0μM [SSB] =0.2nM | [RecA] =2.0μM [SSB] =0.35nM | [RecA] =2.0μM [SSB] =0.65nM | [RecA] =2.0μM [SSB] =1.2nM | [RecA] =2.0μM [SSB] =2.2nM | [RecA] =2.0μM [SSB] =4.2nM | [RecA] =2.0μM [SSB] =7.8nM | [RecA] =2.0μM [SSB] =15nM | [RecA] =2.0μM [SSB] =25nM | [RecA] =2.0μM [SSB] =50nM |
| **H** | [RecA] =2.6μM [SSB] =0 | [RecA] =2.6μM [SSB] =0.1nM | [RecA] =2.6μM [SSB] =0.2nM | [RecA] =2.6μM [SSB] =0.35nM | [RecA] =2.6μM [SSB] =0.65nM | [RecA] =2.6μM [SSB] =1.2nM | [RecA] =2.6μM [SSB] =2.2nM | [RecA] =2.6μM [SSB] =4.2nM | [RecA] =2.6μM [SSB] =7.8nM | [RecA] =2.6μM [SSB] =15nM | [RecA] =2.6μM [SSB] =25nM | [RecA] =2.6μM [SSB] =50nM |

**Table S11. TM3/4 variant and drug concentrations in each well**

| | 1 | 2 | 3 | 4 | 5 | 6 | 7 | 8 | 9 | 10 | 11 | 12 |
|---|---|---|---|---|---|---|---|---|---|---|---|---|
| **A** | WT + 5 μM Lumacaftor | WT + 10 μM Lumacaftor | WT + 100 μM Lumacaftor | WT + 200 μM Lumacaftor | WT + 300 μM Lumacaftor | WT + 400 μM Lumacaftor | WT + 500 μM Lumacaftor | WT + 600 μM Lumacaftor | WT + 800 μM Lumacaftor | WT + 1000 μM Lumacaftor | WT + 1200 μM Lumacaftor | WT + 1600 μM Lumacaftor |
| **B** | V232D + 5 μM Lumacaftor | V232D + 10 μM Lumacaftor | V232D + 100 μM Lumacaftor | V232D + 200 μM Lumacaftor | V232D + 300 μM Lumacaftor | V232D + 400 μM Lumacaftor | V232D + 500 μM Lumacaftor | V232D + 600 μM Lumacaftor | V232D + 800 μM Lumacaftor | V232D + 1000 μM Lumacaftor | V232D + 1200 μM Lumacaftor | V232D + 1600 μM Lumacaftor |
| **C** | E217G + 5 μM Lumacaftor | E217G + 10 μM Lumacaftor | E217G + 100 μM Lumacaftor | E217G + 200 μM Lumacaftor | E217G + 300 μM Lumacaftor | E217G + 400 μM Lumacaftor | E217G + 500 μM Lumacaftor | E217G + 600 μM Lumacaftor | E217G + 800 μM Lumacaftor | E217G + 1000 μM Lumacaftor | E217G + 1200 μM Lumacaftor | E217G + 1600 μM Lumacaftor |
| **D** | Q220R + 5 μM Lumacaftor | Q220R + 10 μM Lumacaftor | Q220R + 100 μM Lumacaftor | Q220R + 200 μM Lumacaftor | Q220R + 300 μM Lumacaftor | Q220R + 400 μM Lumacaftor | Q220R + 500 μM Lumacaftor | Q220R + 600 μM Lumacaftor | Q220R + 800 μM Lumacaftor | Q220R + 1000 μM Lumacaftor | Q220R + 1200 μM Lumacaftor | Q220R + 1600 μM Lumacaftor |
| **E** | WT + 5 μM Galicaftor | WT + 10 μM Galicaftor | WT + 100 μM Galicaftor | WT + 200 μM Galicaftor | WT + 300 μM Galicaftor | WT + 400 μM Galicaftor | WT + 500 μM Galicaftor | WT + 600 μM Galicaftor | WT + 800 μM Galicaftor | WT + 1000 μM Galicaftor | WT + 1200 μM Galicaftor | WT + 1600 μM Galicaftor |
| **F** | V232D + 5 μM Galicaftor | V232D + 10 μM Galicaftor | V232D + 100 μM Galicaftor | V232D + 200 μM Galicaftor | V232D + 300 μM Galicaftor | V232D + 400 μM Galicaftor | V232D + 500 μM Galicaftor | V232D + 600 μM Galicaftor | V232D + 800 μM Galicaftor | V232D + 1000 μM Galicaftor | V232D + 1200 μM Galicaftor | V232D + 1600 μM Galicaftor |
| **G** | E217G + 5 μM Galicaftor | E217G + 10 μM Galicaftor | E217G + 100 μM Galicaftor | E217G + 200 μM Galicaftor | E217G + 300 μM Galicaftor | E217G + 400 μM Galicaftor | E217G + 500 μM Galicaftor | E217G + 600 μM Galicaftor | E217G + 800 μM Galicaftor | E217G + 1000 μM Galicaftor | E217G + 1200 μM Galicaftor | E217G + 1600 μM Galicaftor |
| **H** | Q220R + 5 μM Galicaftor | Q220R + 10 μM Galicaftor | Q220R + 100 μM Galicaftor | Q220R + 200 μM Galicaftor | Q220R + 300 μM Galicaftor | Q220R + 400 μM Galicaftor | Q220R + 500 μM Galicaftor | Q220R + 600 μM Galicaftor | Q220R + 800 μM Galicaftor | Q220R + 1000 μM Galicaftor | Q220R + 1200 μM Galicaftor | Q220R + 1600 μM Galicaftor |
